# MEDIATOR SUBUNIT 25 modulates ERFVII-controlled hypoxia responses in Arabidopsis

**DOI:** 10.1101/2024.01.26.577166

**Authors:** Jos H.M. Schippers, Kira von Bongartz, Lisa Laritzki, Stephanie Frohn, Stephanie Frings, Tilo Renziehausen, Frauke Augstein, Katharina Winkels, Katrien Sprangers, Rashmi Sasidharan, Didier Vertommen, Frank Van Breusegem, Sjon Hartman, Gerrit T. S. Beemster, Amna Mhamdi, Joost T. van Dongen, Romy R. Schmidt-Schippers

**Author notes:** contributed equally (Kira von Bongartz, Lisa Laritzki, Stephanie Frohn, Stephanie Frings, Tilo Renziehausen). Corresponding Authors:* Jos H.M. Schippers Department of Molecular Genetics Seed Development Leibniz Institute of Plant Genetics and Crop Plant Research (IPK) Gatersleben 06466 Seeland Germany; Romy R. Schmidt-Schippers Plant Biotechnology Faculty of Biology University of Bielefeld 33615 Bielefeld Germany. The author responsible for distribution of materials integral to the findings presented in this article in accordance with the policy described in the Instructions for Authors (https://academic.oup.com/plcell/pages/General-Instructions) is: Romy Schmidt-Schippers. **Declaration of conflict of interest**: The authors declare no conflict of interest.

## Abstract

Flooding impairs plant growth through oxygen deprivation, which activates plant survival and acclimation responses. Low-oxygen responses are generally associated with activation of group VII ETHYLENE-RESPONSE FACTOR (ERFVII) transcription factors. However, mechanism and molecular components by which ERFVII factors initiate gene expression are not fully elucidated. Here, we show that the Mediator complex subunit *At*MED25 is recruited by RELATED TO APETALA 2.2 (RAP2.2) and RAP2.12 to coordinate gene expression during hypoxia in *Arabidopsis thaliana*.. The *med25* mutants display reduced low-oxygen stress tolerance. *At*MED25 associates with several ERFVII-controlled hypoxia core genes and its loss impairs transcription under hypoxia due to decreasing RNA polymerase II recruitment. Protein complex pulldown assays demonstrate that the Mediator complex built around *At*MED25 is adjusted under low-oxygen conditions. Moreover, during hypoxia, no functional cooperation between *At*MED25 and the two subunits *At*MED8 and *At*MED16 occurs, contrasting previous observations made for other conditions. In addition, *At*MED25 function under hypoxia is independent from ethylene signalling. Finally, a functional conservation at the molecular level was found for the MED25-ERFVII module between *Arabidopsis thaliana* and the monocot *Oryza sativa*, pointing to a potentially universal role of MED25 in enabling ERFVII-dependent transcript responses to hypoxia in plants.

## Introduction

Increased frequencies of heavy rains due to climate change has made flooding one of the most prevalent and severe abiotic stresses threatening agricultural production (Liu et al., 2023). Flooding stress can occur in the form of submergence of the whole plant or waterlogging, which mainly affects the root system. In either way, flooding represents a major detrimental event leading in the long-term to plant death. Since flooding impairs oxygen diffusion, aerobic respiration for providing energy equivalents is hampered or even blocked. To adapt, plants will induce an acclimation response, including activation of anaerobic respiration, a process which is under strict transcriptional control (Bailey-Serres et al., 2012; Renziehausen et al., 2024).

Acclimation to hypoxia in plants is mediated by a multitude of transcription factors (Gasch et al., 2016; Tang et al., 2021; Eysholdt-Derzsó et al., 2023), but group VII ETHYLENE-RESPONSE FACTOR (ERFVII) transcription factors play a key role in transcriptional reprogramming and subsequent metabolic switching (Giuntoli and Perata, 2018). The most prominent ERFVII member is SUBMERGENCE 1A (SUB1A) that has been successfully used to breed submergence tolerant rice varieties (Bailey-Serres et al., 2010). Genes regulated by SUB1A participate in various processes, including growth, scavenging of reactive oxygen species and anaerobic metabolism (Xu et al., 2006; Fukao et al., 2011; Locke et al., 2018). Moreover, SUB1A directly activates two other ERFVII members, ERF66 and ERF67 during submergence, which in turn activate fermentative pathway genes (Lin et al., 2019). In Arabidopsis, the ERFVII family contains five members, which all have been implicated in flooding tolerance, including RELATED TO APETALA 2.2 (RAP2.2; Hinz et al., 2010), RAP2.3 (Papdi et al., 2015), RAP2.12 (Licausi et al., 2011), and HYPOXIA RESPONSIVE ERF 1 (HRE1) and HRE2 (Licausi et al., 2010). ERFVII-controlled genes in both rice and Arabidopsis comprise fermentative genes, such as *PYRUVATE DECARBOXYLASE 1* (*PDC1*) and *ALCOHOL DEHYDROGENASE 1* (*ADH1*) compensating for impaired ATP synthesis under limited oxygen availability (Licausi et al., 2011; Locke et al., 2018). Arabidopsis ERFVII proteins are controlled in their stability by the current cellular oxygen concentration, as they are substrates of the PLANT CYSTEINE OXIDASE (PCO)-dependent branch of the PROTEOLYSIS 6 (PRT6) N-degron pathway (Licausi et al., 2011; White et al., 2017; White et al., 2018). While in rice, ERF66 and ERF67 are also N-degron substrates (Lin et al., 2019), SUB1A is not (Gibbs et al., 2011). Constitutive expressed RAP-type ERFVII factors can be stored at the plasma membrane where they are protected from proteolysis. For instance, RAP2.12 resides under aerobic conditions at the plasma membrane through interaction with ACYL-COA BINDING PROTEIN 1 (ACBP1), but translocates to the nucleus during low-oxygen conditions upon an energy-related lipid signal (Licausi et al., 2011; Schmidt et al., 2018; Zhou et al., 2020). Moreover, RAP2.2, RAP2.12 and to a lesser extend RAP2.3 act redundantly as principle activators of hypoxia-responsive genes through recognition of the HYPOXIA-RESPONSIVE PROMOTER ELEMENT (HRPE) within hypoxia-responsive target promoters (Gasch et al., 2016). In contrast, HRE1 and HRE2 do not seem to interact with the HRPE. In addition, RAP2.3 promotes apical hook formation and is linked to the ethylene signalling pathway, whereby ethylene promotes RAP2.3 nuclear localisation (Kim et al., 2018). Finally, ERFVII protein abundance and action is controlled by additional environmental cues that include, next to ethylene, NO depletion and retrograde signals (Gibbs et al., 2014; Hartman et al., 2019; Barreto et al., 2022). Although it is relatively well known which transcriptional targets ERFVII factors have, and how their turnover is controlled, still, an important challenge is to understand which molecular determinants enable ERFVII proteins to relay low-oxygen-specific signals to the general transcriptional machinery in order to induce gene expression.

To activate transcription, transcription factors must recruit the general transcriptional machinery consisting of RNA polymerase II (RNAPII) and general transcription factors to target promoters. Bridging signals from specific *cis*-acting transcription factors to RNAPII involves coactivators such as the multi-subunit Mediator complex, which is conserved among eukaryotic organisms (Kornberg, 2005). In Arabidopsis, the Mediator core complex consists of 32 subunits forming three distinct modules: Head, Middle, and Tail (Mathur et al., 2011). In addition, the Kinase module composed of CYCLIN-DEPENDENT KINASE 8 (CDK8), *At*MED12, *At*MED13 and CycC (Mathur et al., 2011) is associated with the core complex, of which CDK8 itself has been reported to possess repressor activity but also to positively affect transcription initiation (Gonzalez et al., 2007; Zhu et al., 2014). While the Head module contacts with RNAPII, the Tail module determines gene-specific functions by interacting with *cis*-acting transcription factors (Dotson et al., 2000). Its flexibility in incorporating different subunits allows the Mediator complex to integrate a wide variety of stress signals.

Specific Mediator subunits in plants modulate growth and stress responses, including plant hormones signalling (Ito et al., 2016; An et al., 2017; Wang et al., 2019; Agrawal et al., 2022; Guo et al., 2022), pathogen responses (Li et al., 2018; Suzuki et al., 2022; Zhang et al., 2023), secondary metabolism (Dolan et al., 2017), root growth (Zhang et al., 2018), fatty acid biosynthesis (Kim et al., 2016) and iron uptake (Zhang et al., 2014). The Arabidopsis Mediator subunit 25 (*At*MED25) was initially identified as regulator of flowering time (Cerdan and Chory, 2003; Bäckström et al., 2007), but was later shown to regulate other developmental processes, such as lateral root growth and root hair differentiation (Sundaravelpandian et al., 2013; Raya-Gonzalez et al., 2014). *At*MED25 positively regulates salinity tolerance and pathogen responses, while it negatively affects plant performance under drought stress (Kidd et al., 2009; Elfving et al., 2011; Ou et al., 2011; Çevik et al., 2012; Zhu et al. 2014). The role of *At*MED25 in controlling transcriptional responses to hormones has been extensively studied. *At*MED25 modulates jasmonic acid, abscisic acid and ethylene signalling and is therefore key to plant-stress responses and signalling pathways (Chen et al., 2012, Yang et al., 2014; An et al., 2017). To execute its function, *At*MED25 physically interacts with multiple transcription factors, including MYC2, the master regulator of jasmonate signalling (An et al., 2017; Liu et al., 2019), DREB2A, ABI5 and several AP2/ERF transcription factors (Elfving et al., 2011; Ou et al., 2011; Çevik et al., 2012; Chen et al., 2012), which represent key regulators of stress responses (drought and temperature) potentially involving abscisic acid and ethylene signalling pathways. Intriguingly, *At*MED25 also interacts with the hypoxia-related RAP2.2 (Ou et al., 2011), suggesting a potential role for *At*MED25 in regulating low-oxygen responses.

In this study, we reveal that the group VII ERF transcription factors RAP2.2 and RAP2.12 from Arabidopsis recruit the Mediator complex to activate hypoxia-responsive genes subsequently allowing adaptation to low-oxygen stress. To achieve this, both factors physically interact with the Tail-located subunit *At*MED25. We demonstrate that *At*MED25 conditionally associates with ERFVII-controlled promoters during hypoxia, restricting transcription initiation to low-oxygen stress conditions. Furthermore, we provide evidence that *At*MED25 constitutes a conserved hub in ERFVII-mediated gene expression in the monocot model plant *Oryza sativa*. Remarkably, most of the ERFVII-controlled hypoxia-responsive genes in Arabidopsis are *At*MED25 dependent. However, as their expression is not completely impaired, unknown coactivators are yet to be discovered. Differential employment of the Mediator complex (via *At*MED25) and potentially other coactivators by specific ERFVII factors may enhance the flexibility of gene induction under hypoxia, thereby satisfying context-specific cellular demands during low-oxygen stress.

## Results

### ERFVII factors interact with the Mediator complex subunit *At*MED25

The Arabidopsis ERFVII factor RAP2.2 was previously identified in a high-throughput screening system for transcription factors interacting with *At*MED25 (Ou et al., 2011). To validate interaction between RAP2.2 and *At*MED25, and to determine if other ERFVII members can interact with *At*MED25 as well, targeted protein-protein interaction assays were performed. First, protein-protein interaction studies in yeast using full-length *At*MED25 in combination with all five Arabidopsis ERFVII transcription factors, namely RAP2.2, RAP2.12, RAP2.3, HRE1 and HRE2 were conducted (**Figure 1A**). *At*MED25 interacted with both RAP2.2 and its homolog RAP2.12. Subsequently, *At*MED25-RAP2.2 and *At*MED25-RAP2.12 binding was validated in an *in-vitro* pulldown approach using CFP-tagged ERFVII proteins and flag-tagged AtMED25 protein (**Figure 1B**). *In planta*, *At*MED25-GFP fusion protein exhibited nuclear localisation under both normoxic and hypoxic conditions (**Figure 1C**). Accordingly, in a follow-up bimolecular fluorescence complementation (BiFC) assay with *At*MED25 and the two ERFVII factors, protein complexes were solely detectable in the nucleus (**Figure 1D**). Next, we examined the capability of other mediator subunits besides *At*MED25 to interact with RAP2.2 and RAP2.12. Eight mediator subunits, namely *At*MED8, *At*MED16, *At*MED19, *At*MED21, *At*MED28, *At*MED32, *At*MED34 and *At*MED36 were selected based on their localisation in the Tail or Middle module of the Mediator complex. The Tail module facilitates complex association with stress-related transcription factors (Yang et al., 2014). For several of the tested subunits, specific binding to stress-related transcription factors has been reported previously (Zhang et al., 2014; He et al., 2021). However, none of the tested subunits could bind to RAP2.2 or RAP2.12 (**Supplemental Figure S1**), suggesting *At*MED25 being the sole component physically linking these ERFVII factors to the Mediator complex.

**Figure 1.**
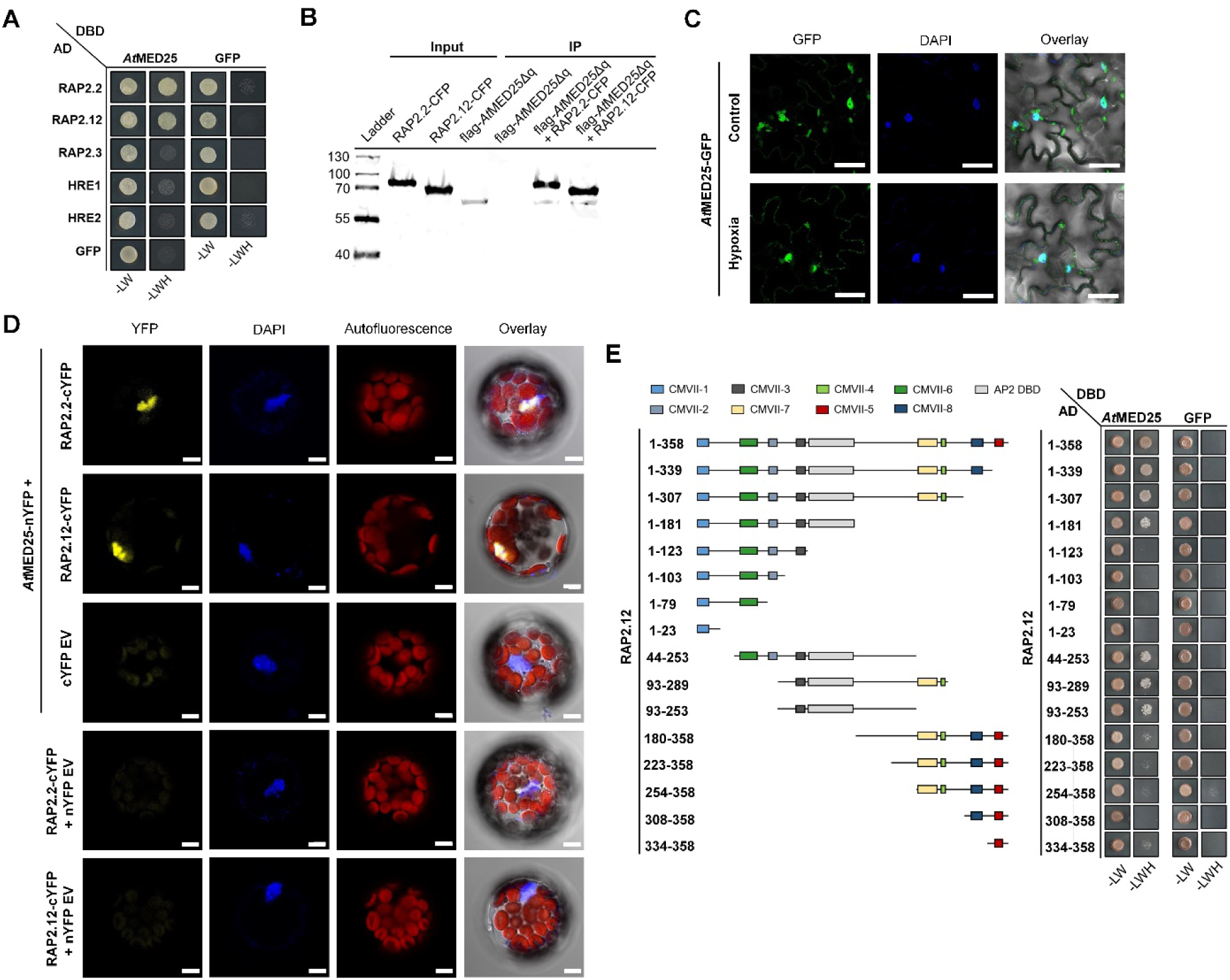
Physical interaction between Mediator subunit *At*MED25 and ERFVII factors. (**A**) Yeast two-hybrid analysis of the interaction between *At*MED25 and five ERFVII factors from Arabidopsis. The *At*MED25 protein was fused with the GAL4 DNA binding domain (DBD), while the ERFVII factors were fused with the GAL4 activation domain (AD). Co-transformed yeast was selected on synthetic dropout (SD) medium lacking leucine and tryptophan (-LW). Protein interactions were tested by transferring the yeast to SD medium lacking leucine, tryptophan and histidine (-LWH) in the presence of 65 mM 3-AT. GFP was used as a negative control. (**B**) *In-vitro* pulldown assay to examine the interaction between *At*MED25 and RAP2.2 or RAP2.12. Purified *in-vitro* expressed proteins were used (flag-*At*MED25Δq, RAP2.2-CFP and RAP2.12-CFP). *At*MED25 was expressed without the glutamine-rich Q-motif to facilitate protein synthesis. GFP affinity beads were used for the pull-down assay. Anti-GFP and anti-flag antibodies were used to detect the respective fusion proteins. As control, 10% of the input samples was loaded. (**C**) Nuclear localisation of *At*MED25-GFP in leaves of stable transgenic Arabidopsis plants under normoxia and after 3 hours of hypoxia, respectively. Nuclei were stained with DAPI. Scale bar: 20 µm. (**D**) BiFC assay of the interaction between *At*MED25 and ERFVIIs. *At*MED25-nYFP and RAP2.2-cYFP or RAP2.12-cYFP or the negative control EV-cYFP constructs were co-transformed into Arabidopsis protoplast to detect protein-protein interaction *in vivo*. YFP fluorescence indicates protein complex formation in nuclei, which were stained with DAPI. As additional negative control, RAP2.2-cYFP and RAP2.12-cYFP were tested against nYFP empty vectors. At least 20 cells were analysed. Scale bar: 5 µm. (**E**) Yeast two-hybrid analysis between *At*MED25 and RAP2.12 to map the interaction domain in RAP2.12. Cloned RAP2.12 fragments are indicated on the left together with the positions of conserved motifs (Nakano et al., 2006). Co-transformed yeast was selected on SD medium lacking -LW. Protein interactions were tested by transferring the yeast to SD medium -LWH in the presence of 65 mM 3-AT. GFP was used as a negative control.

To narrow down regions within RAP2.12 mandatory for *At*MED25 binding, truncated transcription factor proteins lacking individual domains were generated (**Figure 1E**). Several domains conserved in ERFVII factors execute important functions, including oxygen-dependent regulation of protein stability (CMVII-1), DNA binding (AP2 domain), and, in case of RAP2.2 and RAP2.12, transactivation activity (CMVII-5) (Nakano et al., 2006; Bui et al., 2015). In yeast, N-terminus-specific protein fragments of RAP2.12 containing one or multiple conserved motifs upstream of the AP2 domain, i.e., CMVII-1, CMVII-2, CMVII-3 and CMVII-6, failed to bind to *At*MED25. Likewise, C-terminal protein sequences spanning CMVII-4, CMVII-5, CMVII-7 and CMVII-8 were insufficient for interaction. In contrast, the middle part of RAP2.12 (amino acid 93 to 253) containing the AP2 domain with the adjacent upstream domain (CMVII-3) and a downstream flanking sequence was capable of *At*MED25 binding (**Figure 1E**). In conclusion, RAP-specific protein sequences required for *At*MED25 interaction (i.e., the AP2 domain and flanking regions) are non-identical with domains linked to transactivation activity (Bui et al., 2015).

### *At*MED25 confers low-oxygen tolerance through regulating stress-specific transcriptional responses

Physical interference detected between *At*MED25 and the two ERFVII transcription factors, acting as key regulators of transcriptional responses and physiological tolerance towards hypoxia (Licausi et al., 2011; Hinz et al., 2010) suggests involvement of *At*MED25 itself in plant resilience to limited oxygen. Stress tolerance of two independent T-DNA insertion lines lacking a functional *AtMED25* – *med25-1* (*pft1-2*, SALK_129555; Kidd et al., 2009) and *med25-2* (SALK_080230; Xu and Li, 2011) (**Figure 2A**) – was tested by subjecting respective seedlings to anoxia (**Figure 2B-C**). Both mutants showed lower survival scores and thus reduced low-oxygen tolerance compared to the stressed wildtype. Under non-stress conditions, no difference to wild-type seedlings in terms of growth was visible (**Figure 2B**). However, at the age of six weeks, *med25* plants possessed slightly lower rosette size and dry weight (**Figure 2D, F**). Hence, in order to reliably compare performances of adult mutant and wild-type plants under dark-submergence, stress-specific biomass accumulation relative to control conditions was calculated in addition to absolute plant weight. Both *med25-1* and *med25-2* plants exhibited stress-sensitive phenotypes with stronger reduced relative fresh and dry weight than the submerged wildtype (**Figure 2E-F**), indicating *At*MED25 confers low-oxygen stress tolerance during seedling and vegetative stages.

**Figure 2.**
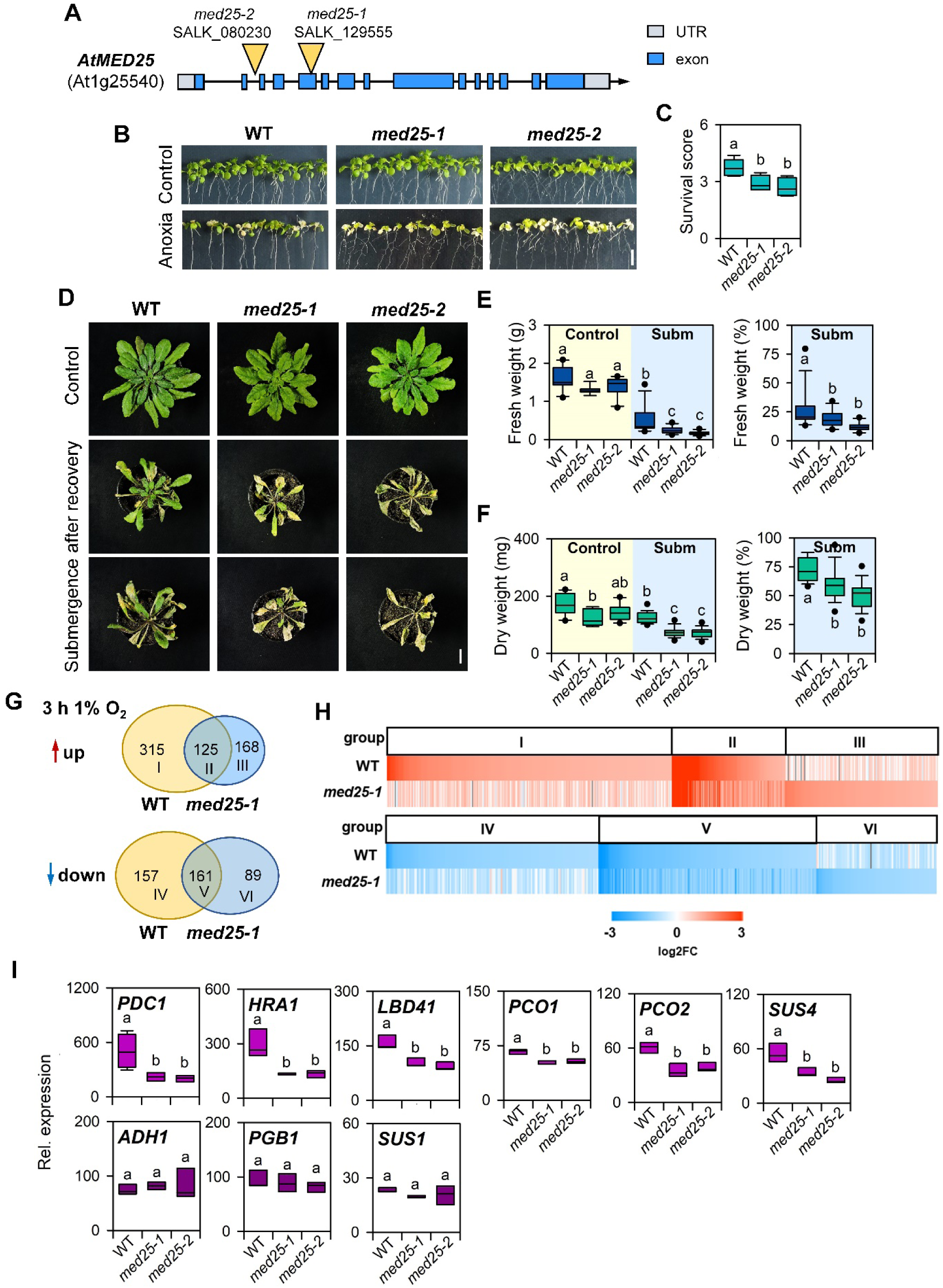
*At*MED25 is a positive regulator of low-oxygen stress adaptation. (**A**) Schematic representation of T-DNA insertion sites *med25-1* (SALK_129555) and *med25-2* (SALK_08230) in *AtMED25*. Location of T-DNA site is indicated with an orange arrow. (**B**) followed by 3 d of recovery. Scale bar: 1 cm. (**C**) Survival scores of *med25* mutants and WT seedlings after recovering from the anoxia treatment. n = 5 (with each 15 seedlings/genotype). Different letters above boxes indicate significant difference to WT according to one-way ANOVA followed by the post hoc Tukey test (*p*<0.05). (**D**) Phenotypes of 6-week-old *med25-1* and *med25-2* and the WT after submergence. Scale bar: 2 cm. (**E** and **F**) Fresh weight (**E**) and dry weight (**F**) of WT, *med25-1* and *med25-2* plants (given as absolute and relative values) under control conditions (n = 8) or after exposure to dark submergence for 3 d followed by 5 d reoxygenation (n = 19). Different letters above bars indicate significantly different groups as determined by one-way ANOVA followed by the post hoc Tukey test (*p*<0.05). (**G**) Venn diagrams display the results of a RNA-Seq analysis for WT and *med25-1* plants under hypoxic conditions. Up- and downregulated responses are represented separately. (**H**) Heatmaps indicate clustered and grouped (I-VI) up- or downregulated genes upon hypoxia (3 h with 1 % O2) in the WT and *med25-1*. Data for the heatmaps are given in **Supplementary Dataset S1 and S2**). (**I**) HCG expression after 3 h of hypoxia treatment in the WT, *med25-1* and *med25-2*. Boxes indicate relative fold change (FC) in expression as compared to normoxic samples (n = 5), as determined by qRT-qPCR. Different letters above boxes indicate significantly different expressed genes according to one-way ANOVA followed by the post hoc Tukey test (*p*<0.05).

As *At*MED25 interacts with ERFVII factors involved in transcriptional reprogramming under low-oxygen stress, we hypothesised that *AtMED25* might affect transcriptional responses to hypoxia. To test this hypothesis, we performed RNA-seq analysis with *med25-1* and wild-type seedlings under normoxic and short-term hypoxic conditions (3 hours, 1 % O_2_). Under normoxic conditions, 2,042 differentially expressed genes (DEGs, FDR ≤ 0.05 and log2FC ≥ 1 or ≤ -1) were identified, as indicated in **Supplemental Dataset S1**. A gene ontology (GO) enrichment analysis revealed that genes involved in the response to stress, especially biotic stress, were upregulated in *med25-1* under control conditions (**Supplemental Dataset S1**). This observation aligns with a reported increased basal pathogen tolerance of *med25* mutants in rice and Arabidopsis (Suzuki et al., 2022; Blomberg et al., 2023). Among the downregulated DEGs, mainly genes involved in cell wall-related processes were present, suggesting altered development or cell wall composition. Upon hypoxia, 758 DEGs were identified for the wildtype, while in *med25-1,* 543 DEGs were found (**Supplemental Dataset S1**). In the wildtype, 440 genes were induced, while in *med25-1* 293 genes responded, indicating for the latter an overall weaker transcriptional response to hypoxia. GO enrichment for genes significantly responding in the wildtype (group I) but not in *med25-1* upon hypoxia mainly related to biotic stress, drought, wounding and ABA (**Supplemental Dataset S2**). Hypoxia core genes (HCGs) are characterised by their strong upregulation under hypoxia (Mustroph et al., 2009), and these were mainly found amongst the overlapping 125 upregulated genes (group II, **Figure 2G**, **Supplemental Dataset S2**). However, in general, HCGs responded weaker in the *med25-1* mutant as compared to the wildtype (**Figure 2H**; **Supplemental Figure S2**). *PDC1*, *HRA1*, *PCO1*, *PCO2*, *HUP9* and *HRE2* were amongst HCGs being lower expressed in *med25-1*, while *PGB1* and *ETR2* showed similar expression levels as in wildtype. Genes only responding in *med25-1* upon hypoxia (group III) had GO enrichment for trehalose metabolism. Not only were less genes induced in *med25-1* as compared to wildtype, also less were downregulated (220 versus 318, respectively) (**Figure 2G, H**). Group IV represents genes significantly downregulated in the wildtype under hypoxia, which enrich for membrane proteins (**Supplemental Dataset S2**). Commonly downregulated genes (group V) comprised genes regulating cell wall organisation and biogenesis. In addition, 89 genes (group VI) were only downregulated in stressed *med25-1*, however, no significantly enriched GO term was obtained. Taken together, *At*MED25 has a profound effect on the transcriptional response to low oxygen in Arabidopsis.

### Induction of a subset of ERFVII target genes under hypoxia is *At*MED25-dependent

The ERFVII transcription factors RAP2.2 and RAP2.12 are the principle regulators of HCGs (Gasch et al., 2016) and we show that they interact with *At*MED25. Next, we validated reduced HCG induction in *med25-1* and *med25-2* plants upon hypoxia using RT-qPCR (**Figure 2I**). While a 3-hour exposure to 1% oxygen resulted in strong induction of ERFVII-controlled hypoxia-responsive genes in wild-type plants, induction was compromised for the genes *PDC1*, *HRA1*, *LBD41*, *PCO1*, *PCO2* and *SUS4* in both *med25* mutants (**Figure 2I**). Notably, in contrast to above mentioned genes, *ADH1* and *PGB1*, but also *SUS1* remained unaffected in their responses to hypoxia in *med25* plants (**Figure 2I**), indicating ERFVII-regulated HCGs group into *At*MED25-dependent and –independent genes.

Physical interaction of *At*MED25 with RAP2.2 and RAP2.12 (**Figure 1A-B** and **D**) and the role of *At*MED25 in regulating ERFVII-dependent gene expression and physiological low-oxygen tolerance (**Figure 2B-F and I**) let us test the transcriptional activity of RAP2.2 and RAP2.12 in the presence and absence of *At*MED25, respectively. For this, in addition to wildtype, transactivation assays were performed in *med25-1* and *med25-2* knock-out backgrounds (**Figure 3A-E**). Induction of luminescence-based reporter constructs including the promoters of *PDC1* (**Figure 3A**), *HYPOXIA-RESPONSE ATTENUATOR1* (*HRA1*, **Figure 3B**) and *STEAROYL-ACYL CARRIER PROTEIN Δ9-DESATURASE6* (*SAD6*) (**Figure 3C**) by RAP2.2 and RAP2.12 was compromised in the absence of *AtMED25* and increased upon *AtMED25* co-expression, respectively. In contrast, induction of *pADH1:LUC* or *pPGB1:LUC* reporters by RAP2.2 and RAP2.12 in *med25* protoplasts was unaffected upon loss of *AtMED25* (**Figure 3D-E**), being in line with *ADH1* and *PGB1* expression responding to hypoxia in *med25* knock-out plants similar to the stressed wildtype (**Figure 2I**). Taken together, *At*MED25 is required for regulation of a distinct set of ERFVII-controlled genes.

**Figure 3.**
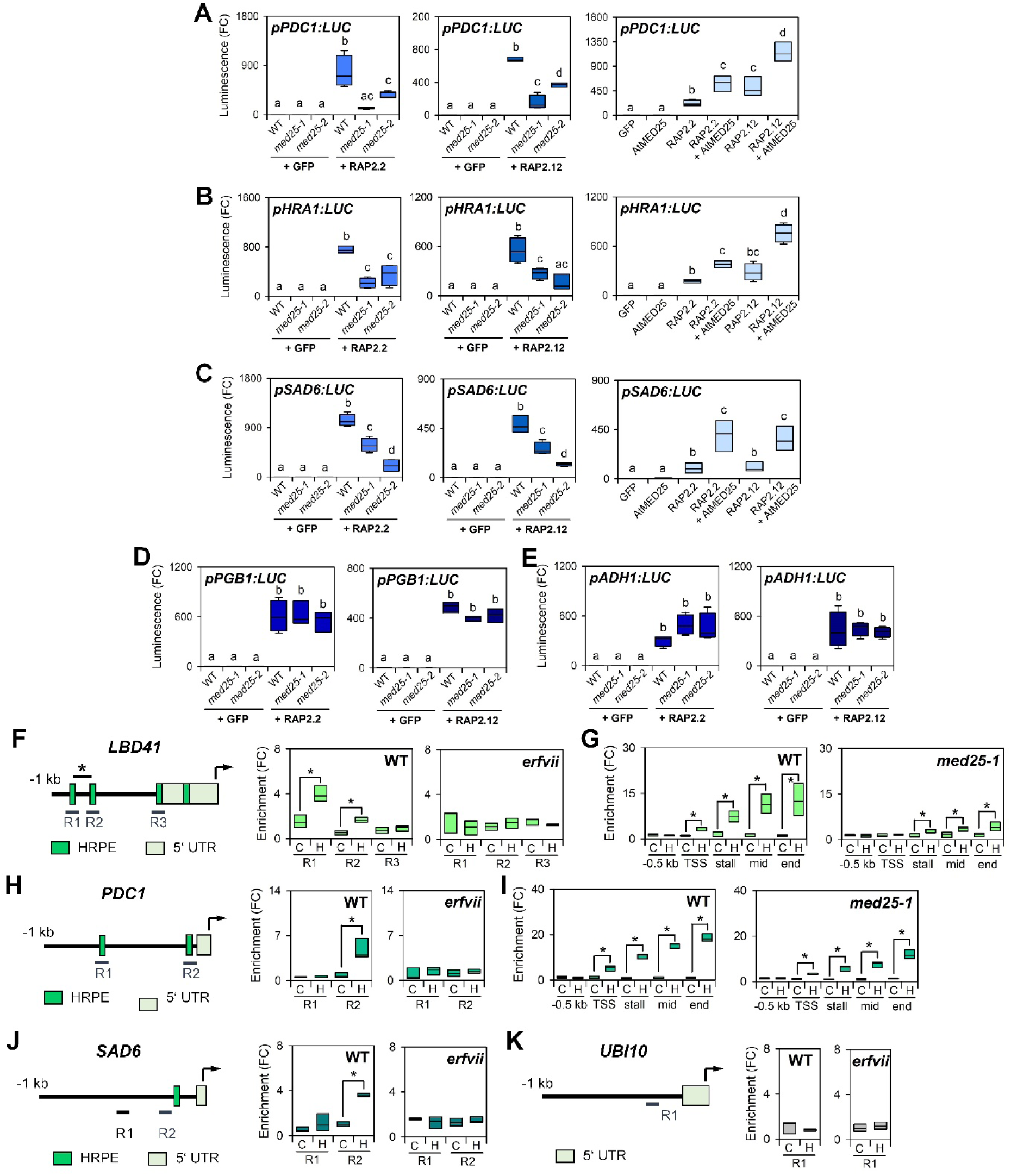
*At*MED25 modulates ERFVII-dependent activation of hypoxia-responsive promoters. (**A-C**) Activity of RAP2.2 and RAP2.12 on (**A**) *pPDC1:LUC*, (**B**) *pHRA1:LUC* and (**C**) *pSAD6:LUC* in protoplasts of wildtype (WT), *med25-1* and *med25-2* (left) and in WT protoplasts upon *AtMED25* co-expression (right). For each promoter and transcription factor, four independent transformation events were analysed. Different letters above bars indicate significantly different groups as determined by one-way ANOVA followed by post-hoc Tukey test (*p*<0.05). A GFP construct was used as negative control. (**D-E**) Unaffected activity of RAP2.2 and RAP2.12 on (**D**) *pPGB1:LUC* and (**E**) *pADH1:LUC* in protoplasts of WT, *med25-*events were analysed. Different letters above bars indicate significantly different groups as determined by one-way ANOVA followed by post-hoc Tukey test (*p*<0.05). A GFP construct was used as negative control (**F, H, J, K**) ChIP assays with *At*MED25-GFP to assess *At*MED25 association with hypoxia-responsive promoters in the WT and *erfvii* background. (**F**) *LBD41*, (**H**) *PDC1*, (**J**) *SAD6*, and (**K**) *UBI10*, the latter serving as negative control. Schemes indicate the upstream genomic regions (R), and the presence of the HRPE motif (green box). Position of the *LBD41* HRPE motif, shown to be bound by RAP2.2 and RAP2.12 (Gasch et al., 2016) is indicated. n = 3. Asterisks in the graphs indicate significant enrichment according to one-way ANOVA followed by post-hoc Tukey test (*p*<0.05). (**G** and **I**) RNAP II recruitment to *LBD41* and *PDC1* promoters in WT and *med25-1* background under control and hypoxic conditions. Map of primer pair locations along the two genes is provided in **Supplemental Figure S3**. ChIP enrichment at each location is referenced to negative control samples (-antibody controls). Values were calculated using the ΔΔCT method. n = 3. Asterisks in the graphs indicate significant enrichment according to one-way ANOVA followed by post-hoc Tukey test (*p*<0.05).

### *At*MED25 association with ERFVII-regulated promoters requires hypoxia

In order to investigate the dynamics of *At*MED25 association with ERFVII-regulated promoters, chromatin immunoprecipitation quantitative PCR (ChIP-qPCR) was conducted using plants expressing GFP-tagged *At*MED25 as bait (**Figure 3F-J**). Next to aerobic conditions, *At*MED25-promoter interactions were studied after 3 hours of hypoxia, which is the moment where ERFVII factors such as RAP2.12 are detectable in the nucleus (Kosmacz et al., 2015). Enrichment of genomic DNA spanning different promoter regions of the tested hypoxia-inducible genes (*PDC1*, *LBD41* and *SAD6*) was analysed. These promoter regions were selected because they contain or are closely located to an HRPE copy previously shown to be recognised by RAP2.2 and RAP2.12 (Gasch et al., 2016). Significant enrichment of *At*MED25 on promoter regions containing HRPEs within *LBD41*, (**Figure 3F**) *PDC1* (**Figure 3H**) and *SAD6* (**Figure 3J**) occurred under hypoxia but not under aerobic conditions, while for the negative control *UBIQUITIN10* (*UBI10*) no *At*MED25 enrichment was detectable under both conditions (**Figure 3K**). To validate *At*MED25 association with hypoxia-responsive promoters is executed through ERFVII factors, the *erfvii* quintuple mutant (Marín-de la Rosa et al., 2014) was included in our analyses. Unlike in the wildtype (**Figure 3F-J**), *At*MED25 failed to bind to hypoxic promoters in *erfvii* upon exposure to hypoxia (**Figure 3F-J**).

To determine whether *At*MED25 is required for recruitment of RNA POLYMERASE II (RNAPII) to hypoxia-inducible, ERFVII-controlled genes, ChIP-qPCR assays were performed with wildtype and *med25-1* plants grown under normoxic and hypoxic conditions using a RNAPII-specific antibody. Subsequently, enrichment of genomic DNA at specific regions along two hypoxia-related genes was tested. As previously described (Hemsley et al., 2014), the following regions were probed: (1) 500 bp upstream of the TSS (-500), (2) the TSS site itself, (3) 50 bp after the start codon (the site of RNAPII stalling, if it occurs), and (4) the middle region of the ORF and 3’UTR region (**Supplemental Figure S3**). Under normoxic conditions, *LBD41* and *PDC1* were transcriptionally nearly undetectable and no significant enrichment for RNAPII along the genomic region was identified (**Figure 3G and I**). However, upon hypoxia, a significant enrichment for RNAPII was observed in the wildtype for both *LBD41* and *PDC1* (**Figure 3G and I**). In comparison to wildtype, considerably weaker enrichment for RNAPII occurred in *med25-1*, showing that *At*MED25 presence is mandatory for optimal RNAPII recruitment toward distinct hypoxia-responsive genes controlled by ERFVIIs.

### *At*MED25 interactome is altered under oxygen constraints

The Mediator complex appears in organisms of several kingdoms as highly flexible structure able to adjust dynamically to environmental signals in terms of its subunit composition, abundance and stability (Dolan et al., 2017). Also, the ability of individual subunits to specifically interact with other Mediator complex components or proteins with different functions, e.g. transcription factors, partially depends on environmental stimuli (He et al., 2021). We investigated if oxygen shortage affects the *At*MED25 interactome to reveal hints towards the response of the Mediator complex as a whole to hypoxia. To test, we performed an immunoprecipitation using GFP-tagged *At*MED25 in Arabidopsis and conducted mass spectrometry analysis (IP-MS) to reveal the specific interactome. Specifically, *35S:AtMED25-GFP* plants were stressed for 3 hours (1% O2) and the *At*MED25-interactome herein was compared with that of plants under non-stress conditions (**Figure 4A-C, Supplemental Dataset S3**). Under control and stress conditions, a multitude of co-purified proteins were identified. 47 of these proteins were also found to be enriched in IP samples from plants subjected to hypoxia. These co-purified interacting partners of *At*MED25 form a complex irrespective of oxygen availability. Among those constitutively present proteins were 16 Mediator subunits, such as *At*MED8, *At*MED11, *At*MED18, *At*MED16, and *At*MED32 belonging to the Head and Tail modules, respectively, as well as *At*MED12, *At*MED13 and CDKE belonging to the associated kinase module. Additionally, proteins with other molecular functions, including two *At*MED25-regulating E3 ubiquitin-protein ligases (MBR1 and MBR2; Iñigo et al., 2012) and two ELKS/Rab6-interacting/CAST family proteins (ELKS1 and ELKS2) were detected. Solely under normoxia, 32 proteins were co-purified with *At*MED25, including four Mediator subunits (*At*MED3, *At*MED9, *At*MED22A and *At*MED30), the kinases ARABIDOPSIS EL1-LIKE 1 (AEL1) and AEL3 and the ubiquitin-binding protein TOM1-LIKE 6 (TOL6). On the other hand, under hypoxia, 65 proteins co-purified with *At*MED25, of which 18 proteins were only enriched under low-oxygen conditions. Hypoxia-specific proteins included, next to the Mediator subunit *At*MED21, also the RNA-binding protein ATRBP45C, the protein kinase RAF20 and the CCCH zinc-finger protein C3H44. Taken together, hypoxia affects the *At*MED25-interactome regarding other Mediator subunits but also functionally diverse proteins, suggesting that protein-protein interaction with *At*MED25, but also the Mediator complex itself is modulated by hypoxia.

**Figure 4.**
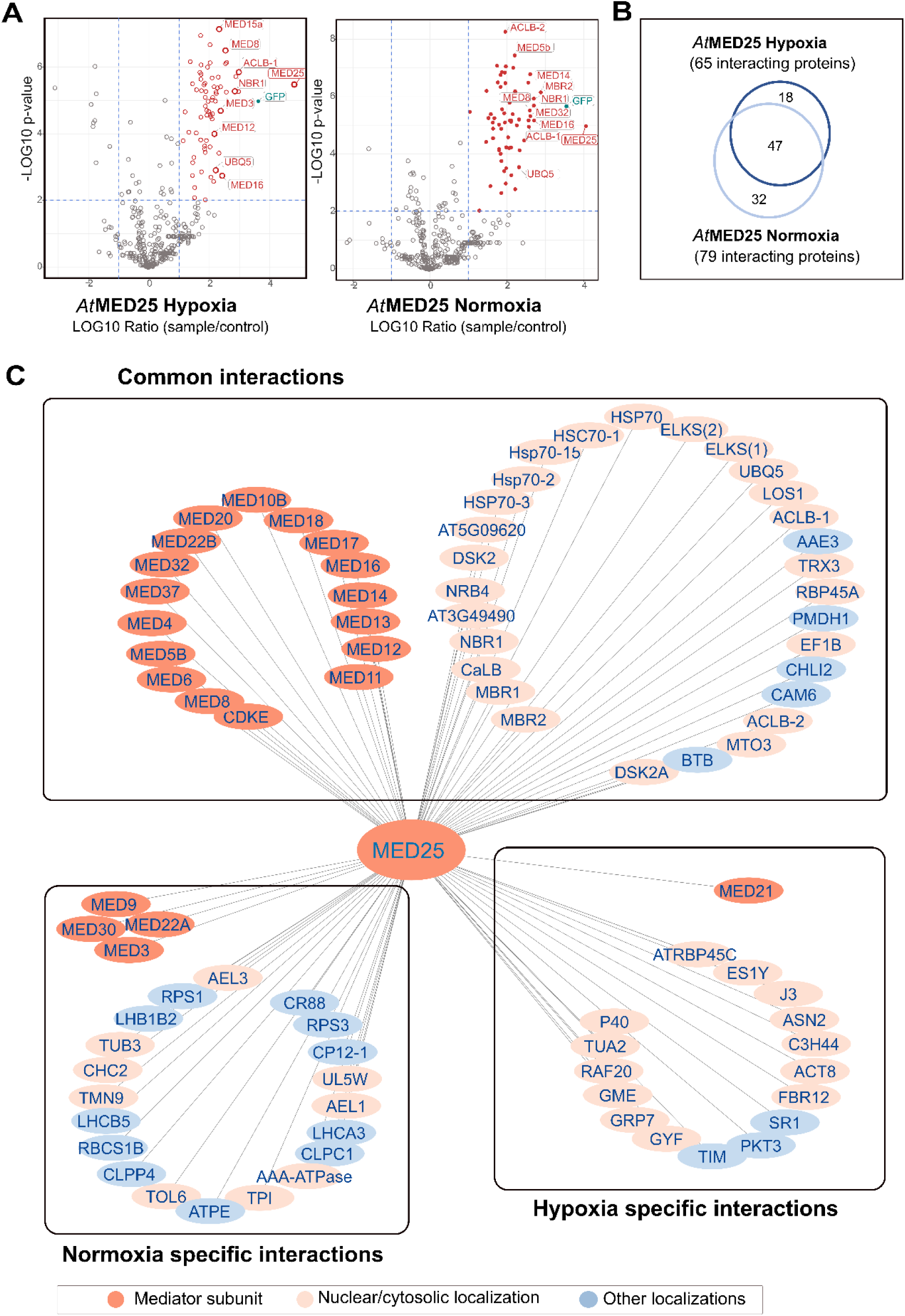
*At*MED25 interactome identified by IP-MS is conditional. (**A**) Volcano plots displaying all identified proteins under normoxia and hypoxia. Proteins depicted in red colour are significantly and highly enriched in the *At*MED25-GFP overexpressors relative to the control samples. (**B**) Venn diagram showing the overlap in the identified interacting proteins under hypoxic and normoxic conditions. (**C**) Interactome of *At*MED25 in the different conditions. The *At*MED25 network was constructed with the Cytoscape software. The different pink (bold or light) to blue colors indicate Mediator complex subunits, proteins localised in the nucleus/cytosol or elsewhere, respectively.

Among those Mediator subunits constitutively interacting *in-planta* with *At*MED25 were *At*MED8 and *At*MED16 (**Figure 4A-C, Supplemental Dataset S3**). Additionally, *At*MED16-*At*MED25 interaction was verified in yeast (**Supplemental Figure S4**). *At*MED16 acts together with *At*MED25 in the context of transcriptional adjustment to environmental iron deficiency (Yang et al. 2014). Specifically, *At*MED25 and *At*MED16 synergistically initiate transcription mediated by transcription factor ETHYLENE-INSENSITIVE 3 (EIN3). Of note, EIN3 associates directly with *At*MED25 controlling some ethylene-mediated responses, while *At*MED16 is incapable of physically binding EIN3. (Yang et al., 2014). As reported for EIN3, we found that RAP2.2 and RAP2.12 interact with *At*MED25, but not with *At*MED16 (**Supplemental Figure S1**). However, also *At*MED8 was reported to have cooperative functions with *At*MED25 in promoting flowering time in Arabidopsis (Bäckström et al., 2007; Yuan et al., 2023). Because of *At*MED8 and *At*MED16 being part of the *At*MED25-interactome irrespective of oxygen availability (**Figure 4A-C, Supplemental Dataset S3**), they might have roles in ERFVII-dependent transcriptional responses during hypoxia. Of note, due to lack of physical interaction with RAP2.2 or RAP2.12 (**Supplemental Figure S1**), *At*MED8 and *At*MED16 effects on transcription factor action would only be indirect.

### *At*MED25 action under hypoxia does not require *At*MED8 or *At*MED16 and is ethylene-independent

To examine a potential contribution of *At*MED8 and/or *At*MED16 to low-oxygen stress tolerance, performances of *med8* (He et al., 2021) and *med16* (*yid1*, Yang et al., 2014) seedlings subjected to anoxia were studied. No difference in survival was detectable as compared to the wildtype (**Figure 5A-B**). Next to that, a role of *At*MED16 in indirectly controlling ERFVII transcription factors was tested by conducting transactivation assays with RAP2.2 and RAP2.12 using *med16* and wildtype protoplasts, respectively. Additionally, ERFVII activities in the wildtype upon AtMED16 co-expression were analysed (**Figure 5C**). In *med16*, ERFVII activities remained unaffected as compared to the wildtype. Likewise, co-expression of *AtMED16* could not increase transcription factor activity in wild-type protoplasts (**Figure 5C**), which is in contrast to previous observations made for *AtMED25* loss and co-expression, respectively (**Figure 3A-C**). Combined, our data suggest that, different from transcriptional responses as part of non-hypoxic processes, *At*MED25 does not act in concert with either *At*MED8 or *At*MED16 to mediate hypoxic responses involving ERFVII factors. Thus, although complex formation of *At*MED25 with *At*MED8 and *At*MED16 is constituted under hypoxia, a joint action is lacking.

**Figure 5.**
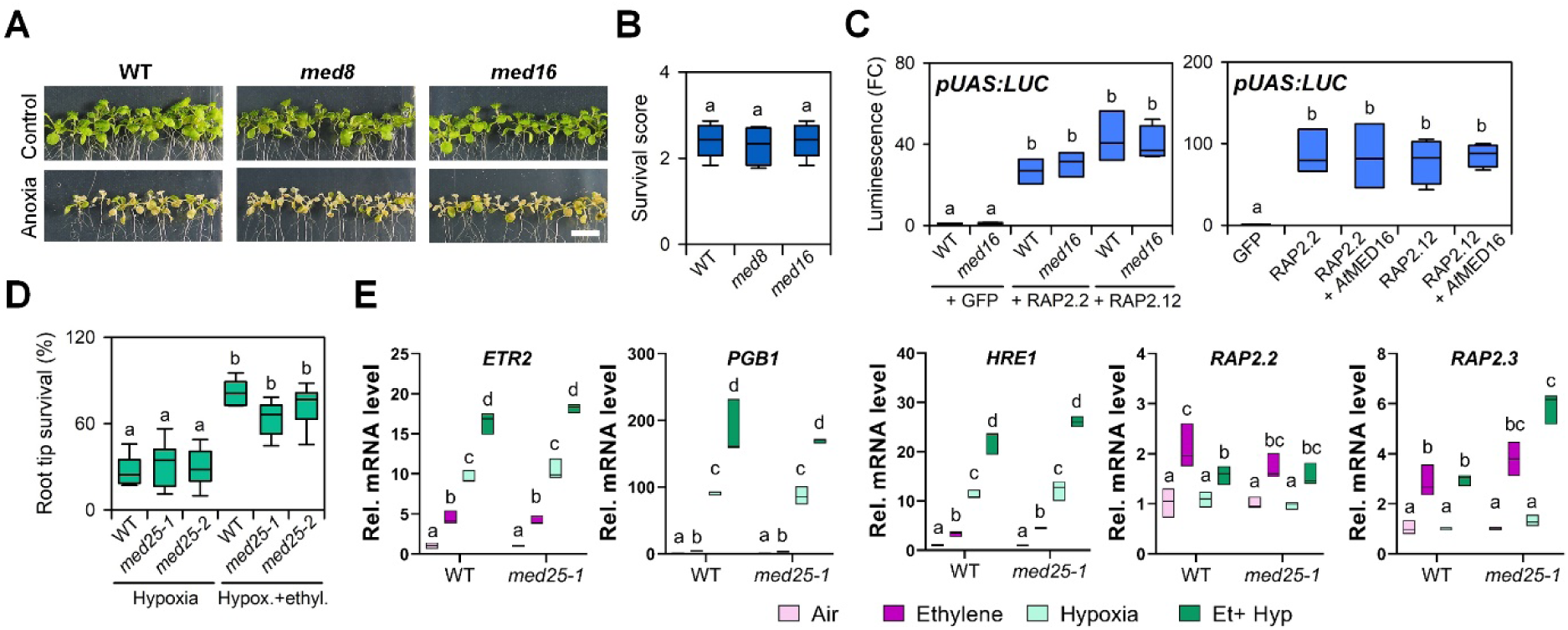
*At*MED25-dependent transcript responses do not involve *At*MED8, *At*MED16 or ethylene. (**A**) Phenotypes of *med8* and *med16* and wildtype (WT) seedlings after 9 h anoxia treatment followed by 3 d of recovery. Scale bar: 1 cm (**B**) Survival scores of *med8* and *med16* mutants and WT seedlings after recovering from anoxia treatment. n = 5 (with each 15 seedlings/genotype). Different letters above boxes indicate significant difference to WT (one-way ANOVA, *p*<0.05). (**C**) RAP2.2 and RAP2.12 activity on *pUAS:LUC* reporter in WT and *med16* protoplasts (left) or in WT protoplasts upon *AtMED26* co-expression (right). For each transcription factor, four independent transformation events were analysed. Different letters above bars indicate significantly different groups as determined by one-way ANOVA followed by post-hoc Tukey test (*p*<0.05). A GFP construct was used as negative control. (**D**) Relative root tip survival of hypoxia-treated (3.5 h) WT, *med25-1* and *med25-2* seedlings (18 rows containing ∼23 seedlings/genotype/treatment) after 4 hours of air/ethylene pre-treatment. Different letters above boxes indicate significant difference to WT (one-way ANOVA, *p*<0.05). (**E**) Relative transcript abundance of ethylene-responsive genes in root tips of WT and *med25-1* seedlings after air/ethylene (4 h) and subsequent hypoxia treatments (4 h). Boxes indicate relative change of mRNA abundance as compared to air-treated control samples (n = 3 with each 2 technical replicates) as determined by RT-qPCR. Different letters above boxes indicate significantly different values (two-way ANOVA followed by post hoc Tukey test (*p*<0.05)).

*At*MED25 (and *At*MED16) action in iron deficiency-related transcriptional responses involves the transcription factor EIN3 acting in an ethylene-dependent manner (Yang et al., 2014). Ethylene is a key phytohormone for plant adaptation to hypoxia through transcriptional, translational and proteolytic control of flooding-induced hypoxia responses (Liu et al., 2022; Cho et al., 2022). Ethylene promotes gene expression and survival under hypoxia partially by enhancing ERFVII stability, which is achieved through blocking internal nitric oxide (NO)- mediated ERFVII proteolysis through increasing the NO-scavenger PGB1 (Hartman et al., 2019). Lack of *At*MED25-*At*MED16 cooperation under hypoxia (**Figure 5A-C**) suggests non-involvement of *At*MED25 in ethylene-mediated responses to hypoxia. Indeed, under oxygen constraints, *med25-1* and *med25-2* mutants benefitted, like the wildtype, from ethylene pre-treatment to improve hypoxia root tip survival (**Figure 5D**). Interestingly, *med25-1* and *med25-2* root hypoxia tolerance was similar to wild-type roots, suggesting that *At*MED25 may control hypoxia tolerance in a shoot-specific manner. Nevertheless, the observed stimulation of mutant performance under hypoxia stress by ethylene indicates limited involvement of *At*MED25 in ethylene-mediated adaptation to hypoxia. In line with this, expression analysis of ethylene-treated *med25-1* and wild-type root tips showed similar induction of the ethylene-responsive genes *ETR2, PGB1, HRE1, RAP2.2* and *RAP2.3* (**Figure 5E**), further supportive of *At*MED25 functioning in ethylene-independent responses to low oxygen.

### The ERFVII-MED25 module is functionally conserved in Arabidopsis and rice

The Mediator complex with its multi-module structure is conserved between eukaryotic organisms (Harper and Taatjes, 2018). Multiple sequence alignment of MED25 proteins from Arabidopsis and several agronomically important crop species, e.g. rapeseed, soy, cotton, wheat, sorghum and rice revealed strong amino acid sequence conservation (**Supplemental Figure S5A-C**). While *At*MED25 is most similar to *Br*MED25 from rapeseed, MED25 proteins from cereal crops form an individual cluster, separating them from dicot species (**Supplemental Figure S5A**). Within the cereal clade, the activator-interacting domain (ACID) of *At*MED25, being sufficient for interaction with stress-induced transcription factors (Ou et al., 2011), is at the amino acid level nearly completely identical (**Supplemental Figure S5B-C**). Hence, we tested whether RAP2.2 and RAP2.12 are capable of interacting with full-length *Os*MED25 from rice, which was indeed the case in yeast (**Figure 6A**). Moreover, the rice ERFVII transcription factor SUB1A, having key regulatory function in hypoxic responses and conferring tolerance to submergence (Xu et al., 2006; Locke et al., 2018), was capable of interacting with *At*MED25 as well as with *Os*MED25 (**Figure 6A**). It suggests that complex formation between ERFVIIs and MED25 proteins may represent a conserved feature in the regulation of low-oxygen responses in multiple plant species. To reveal whether *Os*MED25 acts as positive regulator of ERFVII transcription factor activity from Arabidopsis, we tested RAP2.2 and RAP2.12 activities on different hypoxia-responsive target promoters. Induction of *PDC1*, *HRA1* and *SAD6*-specific luminescence-based reporters by both RAP2.2 and RAP2.12 was increased in the presence of *Os*MED25 (**Figure 6B**). Of note, in all assays performed, *Os*MED25 did not significantly affect target promoter activity in the absence of ERFVII transcription factors. As SUB1A forms nuclear complexes with MED25 proteins in rice protoplasts (**Figure 6C**), the effect of *Os*MED25 on SUB1A performance on reporter constructs derived from the SUB1A target promoters *OsERF66* and *OsERF67* (Lin et al., 2019) was investigated. This revealed significantly increase in SUB1A activity in the presence of *Os*MED25 (**Figure 6D**). These data indicate functional conservation of the MED25-ERFVII module in Arabidopsis and rice at the molecular level. To further demonstrate that MED25 proteins from both species have similar functions in adaptation to low-oxygen stress at whole-plant level, the ability of ectopic *OsMED25* expression to restore wild-type performance in the Arabidopsis *med25-1* mutant was tested. For this, either *35S:OsMED25* or *35S:AtMED25* constructs were introduced into the mutant background and subsequent tolerance assays were performed (**Figure 6E-F**). *med25-1/35S:OsMED25* seedlings subjected to anoxia showed a wildtype-like phenotype, i.e., a higher survival score and thus higher tolerance than observed for *med25-1*, which was a comparable finding as for *35S:AtMED25*-complemented *med25-1* (**Figure 6E-F**). This indicates similar roles of MED25 from both species in establishing physiological resilience toward low-oxygen stress.

**Figure 6.**
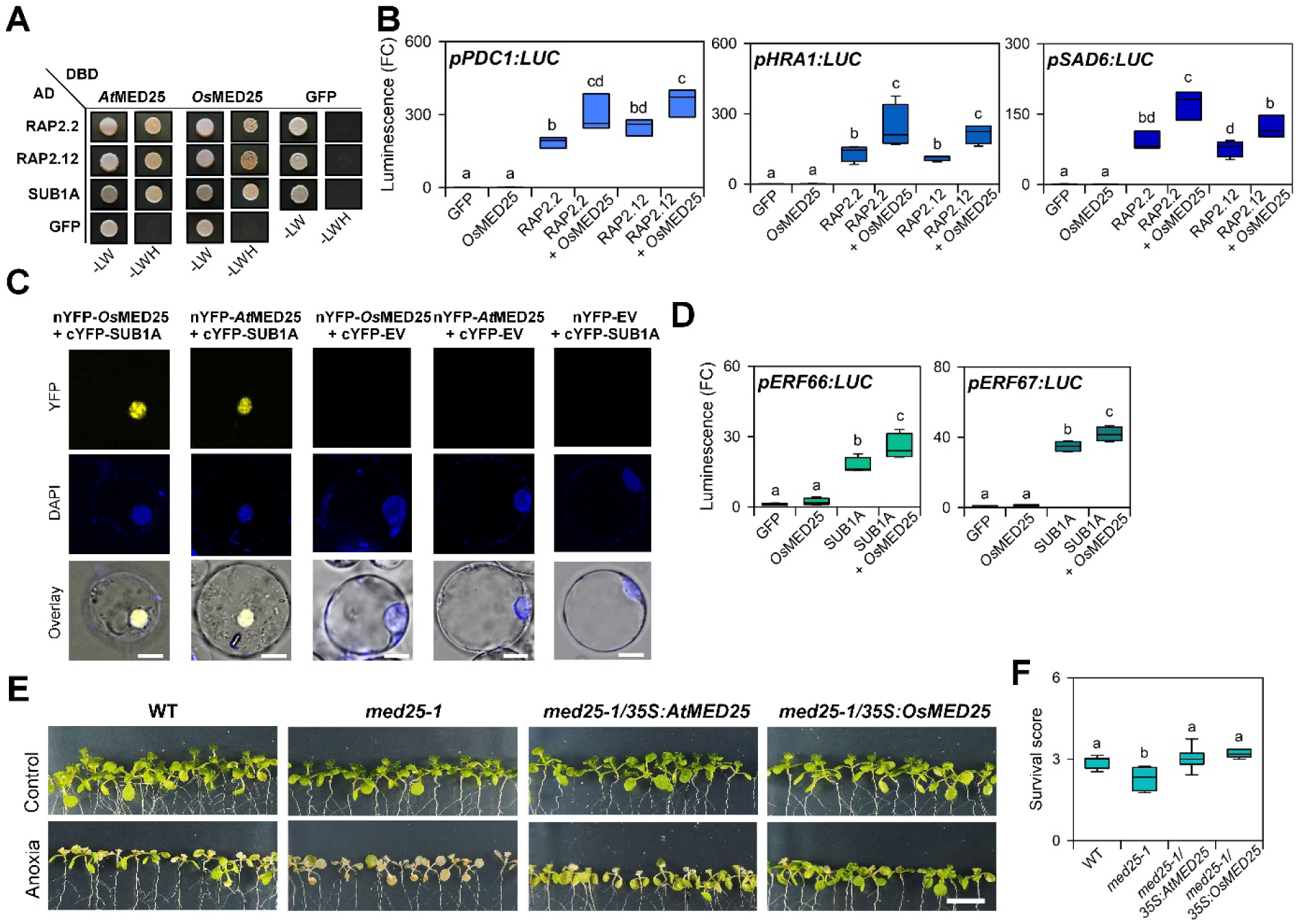
Functional conservation of the ERFVII-MED25 module in Arabidopsis and rice. (**A**) Yeast two-hybrid assay with Arabidopsis and rice ERFVII factors combined with *At*MED25 and *Os*MED25, respectively. Co-transfected yeast was selected on synthetic dropout (SD) medium lacking leucine and tryptophan (-LW). Protein interactions were tested by transferring the yeast to SD medium lacking leucine, tryptophan and histidine (-LWH) in the presence of 65 mM 3-AT. GFP is used as negative control. (**B**) RAP2.2 and RAP2.12 activity on three reporter constructs in Arabidopsis wildtype (WT) protoplasts upon *OsMED25* co-expression was analysed. For each promoter and transcription factor, four independent transformation events were analysed. Different letters above bars indicate significantly different groups as determined by one-way ANOVA followed by post-hoc Tukey test (*p*<0.05). A GFP construct was used as negative control. (**C**) BiFC assay using split-YFP with N-terminus of YFP fused to *Os*MED25 or *At*MED25 and C-terminus to SUB1A. YFP signal indicates protein complex formation in nuclei, which were stained with DAPI (blue). As negative controls, each tested construct was combined with the respective empty vector. Scale bar: 5 µm. (**D**) SUB1A activity on *pERF66:LUC* (left) and *pERF67:LUC* (right) in rice WT protoplasts upon *OsMED25* co-expression. For each promoter four independent transformation events were analysed. Different letters above bars indicate significantly different groups as determined by one-way ANOVA followed by post-hoc Tukey test (*p*<0.05). A GFP construct was used as negative control. (**E**) Phenotypes of WT, *med25-1* and *med25-1* complemented with either *35S:AtMED25* or *35S:OsMED25* after 9 h anoxia treatment followed by 3 d of recovery. Scale bar: 1 cm (**F**) Survival scores of respective genotypes in E after recovering from the anoxia treatment. n = 5 (with each 15 seedlings/genotype). Different letters above boxes indicate significant difference to WT (one-way ANOVA, *p*<0.05).

## Discussion

Plants, as all aerobic organisms, require oxygen for proper functioning and survival by generating respiratory energy. Environmental constraints can limit the amount of oxygen available to sustain plant metabolism and growth, especially during flooding events (Sasidharan et al., 2021). Upon submergence, several acclimation mechanisms are activated to minimize detrimental effects of low oxygen. While ERFVII transcription factors are key regulators of transcriptional reprogramming during hypoxia (Xu et al., 2006; Hinz et al., 2010; Gibbs et al., 2011; Licausi et al., 2011), also other transcription factors have crucial roles in protecting plants from harmful consequences of flooding stress (Liu et al., 2021; Eysholdt-Derzsó et al., 2023). In this study, we show that the ERFVII factors RAP2.2 and RAP2.12 interact with the Mediator complex through subunit *At*MED25 in order to recruit RNAP II to promoters of a subset of hypoxia-responsive genes (**Figure 7**). Furthermore, we demonstrate that *At*MED25 is required for submergence tolerance and that the composition of the Mediator complex attached to *At*MED25 is modulated upon low oxygen stress. In addition, *Os*MED25 from rice is able to interact and promote the transcriptional activity of SUB1A in rice protoplast. These results indicate that the potentially conserved MED25-ERFVII module enables plants to fine-tune their low-oxygen response to promote survival.

**Figure 7.**
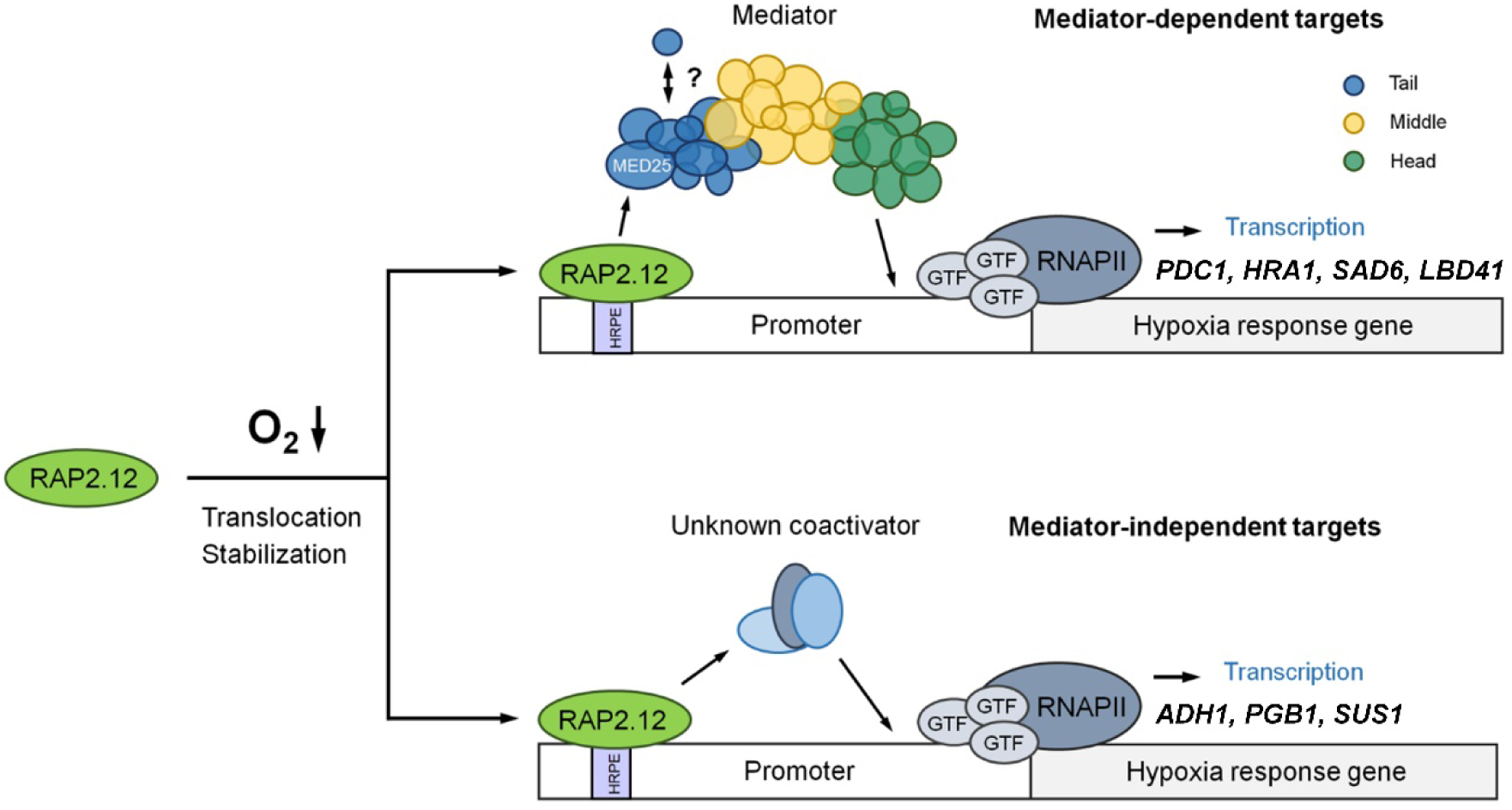
Proposed role of *At*MED25 action in ERFVII-dependent gene regulation under low-oxygen stress. ERFVII transcription factors RAP2.2 and RAP2.12 bind to the HYPOXIA-RESPONSIVE PROMOTER ELEMENT (HRPE) within promoters of target genes such as *PDC1* and *LBD41*. Via their C-termini, both ERFVII factors physically interact with the Tail-specific *At*MED25 subunit as part of the Mediator complex. Association of *At*MED25 with ERFVII target promoters occurs under hypoxia only, likely through low-oxygen induced accumulation of ERFVIIs in the nucleus. The ability of *At*MED25 to interact with other proteins is affected in an oxygen-dependent manner. Initiation of hypoxic gene transcription upon recruitment of RNAPII results in hypoxia tolerance. Next to RAP2.2 and RAP2.12 from Arabidopsis, interaction of the rice counterpart SUB1A with MED25 proteins from both rice and Arabidopsis and subsequent transcription factor activation was shown, suggesting that the ERFVII-MED25 module is functionally conserved in the plant kingdom.

In Arabidopsis, ERFVII family members have been shown to have partial overlapping and additive functions during submergence (Gasch et al., 2016; Zubrycka et al., 2023), whereby especially RAP2.2, RAP2.3 and RAP2.12 contribute to the expression of HCGs. Here, we found that out of the five ERFVIIs only RAP2.2 and RAP2.12 interact with *At*MED25 (**Figure 1A**). Specifically, RAP2.12 interaction with *At*MED25 is enabled by a transcription factor region containing the DNA-binding domain and its flanking regions, and does not require the well-known C-terminal CMVII-5 domain essential for transcriptional activity (Bui et al., 2015). *At*MED25 interacts with specific members of diverse transcription factor families, including the AP2/ERF family (Ou et la., 2011; Çevik et la., 2012), bHLH members (Çevik et la., 2012), bZIP proteins (Chen et al., 2012), MYB family members (Çevik et la., 2012), and several AUXIN RESPONSE FACTOR (ARF) proteins (Ito et al., 2016). As observed in our study, also in DREB2A the protein domain required for *At*MED25 interaction does not correspond to the transcriptional activation domain (Elfving et al., 2011). This suggests that separation of the *At*MED25-interaction domain from the transcriptional activation domain might be a common feature in AP2/ERF proteins. In contrast, the VIRUS PROTEIN 16 (VP16) transcriptional activation domain directly interacts with the ACID domain of human MED25 (Vojnic et al., 2011), indicating that *At*MED25 binds to some proteins directly via their transcriptional activation domains. Improved understanding of the interaction modes between *At*MED25 and the diverse transcription factors will greatly advance our understanding of transcriptional regulation in plants.

The Mediator complex acts as an information hub that conveys signals from transcription factors to RNAP II to modulate and regulate gene expression to guide development and secure plant survival (Agrawal et al., 2021). Especially *At*MED25 appears to act as crucial signalling component, since it is involved in various developmental and stress signalling pathway (Kazan, 2017; Zhai et al., 2020; Deng et la., 2023). Specific interaction of *At*MED25 with RAP2.2 and RAP2.12 suggests a role of *At*MED25 in low-oxygen tolerance of Arabidopsis. Indeed, we provide evidence that loss of *AtMED25* decreases tolerance to low-oxygen conditions as demonstrated by anoxia and submergence assays (**Figure 2B and D**). Moreover, transcriptional responses to short-term hypoxia were severely affected in *med25* (**Figure 2G and H**). Of note, several hypoxia-induced ERFVII target genes, mandatory for acclimation to low-oxygen stress, showed impaired induction in *med25* plants. These genes comprise *PDC1*, *HRA1*, *HUP7*, *HUP9*, *LBD41*, *PCO1*, *PCO2* and *SUS4* (**Figure 2I**). However, other HCGs like *ADH1*, *PGB1* and *SUS1*, which also contain an ERFVII-specific HRPE element in their promoters (Gasch et al., 2016) remain unaffected in their transcriptional response to hypoxia (**Figure 2I**). As RAP2.3 acts partially redundantly to RAP2.2 and RAP2.12 (Gasch et al., 2016), but does not interact with *At*MED25 (**Figure 1A**), it might explain why some HCGs remain unaffected by the absence of *At*MED25. Alternatively, RAP2.2 and RAP2.12 might also recruit other coactivators in addition to *At*MED25. To determine if the transcriptional activity of RAP2.2 and RAP2.12 relies solely on *At*MED25, we performed transactivation assays in the wild-type and *med25* mutant background using ERFVII target promoter reporters (**Figure 3A-E**). We observed that in the *med25* mutant, RAP2.2 and RAP2.12 function was substantially impaired, but not completely abolished, suggesting additional coactivators exist for RAP2.2 and RAP2.12. A screen for potential interactions with other Mediator components did not reveal additional subunits interacting with RAP2.2 or RAP2.12 (**Supplemental Figure S1**). It would be of interest to identify other coactivators acting in concert with *At*MED25-independent ERFVII factors, i.e., RAP2.3, HRE1 and HRE2. In analogy, the main regulator of transcriptional adaptation to hypoxia in humans, HIF1A, requires the CDK8-Mediator module to regulate a subset of its target genes (Galbraith et al., 2013). However, another subset of HIF1A-regulated hypoxia-responsive genes requires interaction with the TIP60 coactivator complex (Perez-Perri et al., 2016). Recruiting different coactivators may improve the flexibility of HIF1-regulated transcription under hypoxia and thus allows for a context-dependent cellular and phenotypic response. A potentially similar palette of co-activators assisting in ERFVII-dependent transcript regulation may enable flexible adaptation to hypoxia in plants.

To understand if *At*MED25 associates with promoters of hypoxia-responsive genes controlled by ERFVII factors, a ChIP analysis was performed (**Figure 3F-K**). Under control conditions, no enrichment for *At*MED25 at the promoters of *LBD41*, *PDC1* and *SAD6* was observed. However, under hypoxia, *At*MED25 associates with these hypoxia-responsive genes, indicating that binding of *At*MED25 to promoters is conditional. Especially, *At*MED25 was enriched on promoter regions encompassing an HRPE copy, suggesting that its association is ERFVII-dependent. Indeed, simultaneous loss of all five *ERFVII* genes as in the *erfvii* mutant (Marín-de la Rosa et al., 2014) abolished binding of *At*MED25 to promoters under low-oxygen stress (**Figure 3F, H** and **J**). These findings support a conditional relation between ERFVII transactivation activity and *At*MED25 function. In part, this can be explained by stabilisation and translocation of ERFVII factors as a consequence of hypoxia. In this manner, precocious activation of hypoxic responses is prevented. However, it cannot be excluded that also posttranslational modifications and *At*MED25 abundance, as previously shown in the context of flowering time control (Iñigo et al., 2012), might add to the conditional activation of hypoxia-activated genes.

The Mediator complex is highly flexible and its structure and composition are dynamically regulated depending on developmental and environmental cues (Dolan and Chapple, 2017). To obtain insights into the modulation of *At*MED25 function during hypoxia, protein complex pulldown assays were performed under stress and non-stress conditions. These assays identified 16 Mediator subunits belonging to either the Head, Middle or Tail module of the Mediator complex, which constitutively interact with *At*MED25 irrespective of the actual oxygen availability (**Figure 4C**). In contrast, *At*MED21 only associated with *At*MED25 under hypoxia. Notably, *At*MED21 is part of the Middle module of the Mediator complex enabling RNAP II association with the Mediator complex to promote transcription of target genes (Sato et al., 2016; Huang et al., 2021). Moreover, it was shown that *At*MED21 is a target of the co-repressor TOPLESS to dampen transcriptional activation (Leydon et al., 2021). Specific incorporation of *At*MED21 into *At*MED25 protein complexes indicates formation of an active Mediator complex under hypoxia. Potentially, this complex formation might rely on the interaction between ERFVII factors and *At*MED25. However, the inherent low abundance of transcription factor proteins limited their detection in our IP-MS approaches. Next to that, under hypoxia, the chaperone J3 associated with *At*MED25 (**Figure 4C**), which was previously shown to promote heat and drought stress tolerance in Arabidopsis (Wang et al., 2021). Furthermore, the Raf-like MAP kinase kinase kinase RAF20 only interacted with *At*MED25 under hypoxia (**Figure 4C**). RAF20 is known to activate subclass I SUCROSE NONFERMENTING-1-RELATED PROTEIN KINASES 2 (SnRK2s) as part of osmotic stress responses (Soma et al., 2020), which is also an aspect of submergence experience. Taken together, the *At*MED25 interactome composition is modulated in an oxygen-dependent manner. Understanding the molecular mechanisms controlling the configuration of *At*MED25 interactions during its diverse roles in plant development and stress adaptation represents a major challenge for future research.

*At*MED25 reportedly cooperates with *At*MED16 to control EIN3 and EIL1 as part of iron deficiency responses (Yang et al., 2014), while in tomato, MED25 regulates fruit ripening through modulation of ethylene-dependent pathways (Deng et al., 2023). Next to that, *At*MED25 and *At*MED8 are both required for regulating transcriptional changes upon glucose treatment (Seguela-Arnaud et al., 2015). In our study, both *At*MED8 and *At*MED16 were identified as constitutive components of the *At*MED25-specific interactome (**Figure 4C**). Although both subunits do not directly interact with RAP2.2 and RAP2.12 (**Supplemental Figure S1**), their potential contribution to low-oxygen tolerance was examined. In contrast to *At*MED25, *At*MED8 and *At*MED16 are dispensable for anoxia tolerance (**Figure 5A-B**). In line with that, transcriptional activity of RAP2.2 and RAP2.12 towards hypoxia-responsive promoters was not impaired in the *med16* background (**Figure 5C**). As *At*MED25 is linked to the regulation of EIN3 and EIL1 (Deng et al., 2023), it was tested if it also participates in ethylene-mediated adaptation to hypoxia (Liu et al., 2022). Interestingly, *At*MED25 is not mandatory for ethylene-mediated root tip survival (**Figure 5D**), indicating that ethylene-mediated hypoxia acclimation acts through ERFVIIs, but not *At*MED25. In line with this, transcript abundance of genes induced by ethylene was similar in *med25-1* and the wildtype exposed hypoxia (**Figure 5E**).

The *At*MED25 subunit is highly conserved between eukaryotes, suggesting that the observed role in Arabidopsis in hypoxia adaptation may also be important for other plants species. As demonstrated previously, in both *Physcomitrium patens* and Arabidopsis, MED25 promotes salinity tolerance (Elfving et al., 2011). Furthermore, MED25 from wheat complements phenotypic aspects of the Arabidopsis *med25-1* knock-out mutant, including delayed flowering and disease resistance (Kidd et al., 2009). Conservation of MED25 function and amino acid sequence within the plant kingdom suggests a common role in hypoxia signalling in plants (**Supplemental Figure S5**). Here we show that MED25 proteins from Arabidopsis and rice are capable of interacting with RAP2.2, RAP2.12 (both Arabidopsis) and SUB1A (rice) (**Figure 6A**). Furthermore, *Os*MED25 promoted the activity of RAP2.2 and RAP2.12 towards target promoters, whilst it could also stimulate activation of rice *ERF66* and *ERF67* by SUB1A (**Figure 6B** and **D**; Lin et al., 2019). Finally, we demonstrated that *Os*MED25 functionally replaces *At*MED25 *in planta* by restoring low-oxygen tolerance of respectively complemented *med25-1* mutant plants (**Figure 6E-F**). How exactly *Os*MED25 contributes to SUB1A activity will greatly contribute to our knowledge of ERFVII-dependent gene regulation in plants. Our findings add a new facet to the mechanisms by which plants perceive and adapt to low-oxygen stress at the molecular level. Potentially, the provided identification of a conserved role of MED25 in low-oxygen adaptation could help to improve flooding stress resilience in crops.

## Methods

### Plant material

For all indicated experiments, wild-type plants with Columbia-0 background were used. The knock-out mutants *med25-1* (*pft1-2*, SALK_129555; Kidd et al., 2009) and *med25-2* (SALK_080230; Xu and Li, 2011) were described previously and genotyped by PCR with the following primers: 5’-ATGACTTAAGGCAGCACATGC-3’ (LP) and 5’-GTGCTTCTGTCAGCGATTTTC-3’ (RP) for wild-type allele detection in *med25-1*, and 5’-AGGTGTTGGCAATATGTGAG-3’ (LP) and 5’-CAACGCATTCATAAAGCAAT-3’ (RP) for wild-type allele detection in *med25-2*. Detection of mutated alleles was done by PCR with the primer 5’-ATTTTGCCGATTTCGGAAC-3’ combined with RP. The knock-down line *med16* (*yid1*), *med8* mutant (SALK_092406) and *erfvii* mutant were described previously (Yang et al., 2014; Marín-de la Rosa et al., 2014; He et al., 2021).

### Growth conditions and low-oxygen treatments

Tolerance to anoxia was tested for *med25-1*, *med25-2*, *med25-1/35S:AtMED25* and *med25-1/35S:OsMED25* as described previously (Schmidt et al., 2018). Survival scores were calculated as described elsewhere (Gibbs et al., 2011). Description of conducted submergence assays with adult *med25-1* and *med25-2* plants can be found elsewhere (Schmidt et al., 2018). For ethylene and hypoxia treatments, root tip tolerance assays and RT-qPCR on *in vitro* grown seedlings on agar plates, the same conditions and methods were used a reported previously (Hartman et al., 2019; Liu et al., 2022).

### Cloning of constructs

For interaction studies in yeast, full-length *AtMED25* and *OsMED25* coding sequences in pENTR-D/TOPO (Invitrogen) were together with coding sequences of *AtMED8*, *AtMED16*, *AtMED19*, *AtMED21*, *AtMED28*, *AtMED32*, *AtMED34* and *AtMED36* (kindly provided in pENTR-D/TOPO by Prof. Li-Jia Qu, School of Life Sciences, Peking University, Beijing, China) recombined into pDEST32 (Thermo Fisher Scientific, Waltham, USA), possessing the GAL4 DNA-binding domain, as bait constructs. *AtMED25* coding sequence, *RAP2.2* and *RAP2.12* full-length and truncated coding sequences as well as the *SUB1A* coding sequence were cloned into pENTR-D/TOPO and together with *HRE1*, *HRE2* and *RAP2*.3 in pENTR-D/TOPO (kindly provided by Prof. Margret Sauter, Christian-Albrechts University, Kiel, Germany) recombined into pDEST22 (Thermo Fisher Scientific, Waltham, USA), possessing the GAL4 activation domain, as prey constructs.

Transactivation assays were conducted using *PDC1*, *SAD6*, *HRA1*, *ADH1* and *PGB1* promoters recombined into p2GWL7,0 and *RAP2.12* and *RAP2.2* coding sequences into p2GW7,0 (Bui et al., 2015; Karimi et al., 2002).

For bimolecular fluorescence complementation analysis, *MED25* coding sequences from Arabidopsis and rice were recombined into pDH51-GW-YFPn, while full-length *RAP2.12*, *RAP2*.*12* and *SUB1A* coding sequences in pENTR-D/TOPO (Invitrogen) were recombined into pDH51-GW-YFPc (Zhong et al., 2008).

For *in vitro* pulldown experiments the coding region of *At*MED25Δq was fused N-terminally with a flag-tag by PCR with oligonucleotides containing restriction site overhangs for cloning into pF3A WG BYDV (Promega). RAP2.12 C-terminally fused to CFP was previously established (Schmidt et al., 2018). Similarly, the CDS of RAP2.12 was C-terminally fused to CFP by PCR and cloned into pF3A WG BYDV.

All oligonucleotide sequences used for cloning of constructs are listed in **Supplemental Table S1**.

### Plant transformation

Full-length *AtMED25* and *OsMED25* coding sequences and the full-length *AtMED25* promoter were cloned into pENTR-D/TOPO (Invitrogen) and recombined either into pK7WG2 (*AtMED25* and *OsMED25* coding sequences) (Karimi et al., 2002) or pK7FWG2 (only *AtMED25*) providing the 35S promoter (Karimi et al., 2002) or pKGWFS7 (*AtMED25* promoter) (Karimi et al., 2002). After transformation into Agrobacterium, *35S:AtMED25* and *35S:OsMED25* were transformed into *med25-1*, *35S:AtMED25-GFP* into wildtype and *erfvii* and *pAtMED25:GUS* into wild-type plants using the floral dip method. Antibiotic-resistant T1 seedlings were identified and, in case of AtMED25-GFP, tested for a present GFP signal by confocal microscopy analysis.

### Interaction studies in yeast

Recombinant pDEST22 and pDEST32 vectors were co-transformed into the *Saccharomyces cerevisiae* strain PJ694a (James et al., 1996) using the LiAc method and successfully transformed yeast colonies were selected on solid SD medium lacking Trp and Leu. Subsequently, single colonies were grown overnight (30°C, 300 rpm) on selective SD medium and 3 µl were spotted on SD medium lacking Trp, Leu and His and containing 65 mM 3-amino-1,2,4-triazole. As negative controls, recombinant pDEST22 or pDEST32 vectors were co-transformed with the corresponding counterpart containing GFP.

### Transactivation assays and bimolecular fluorescence complementation analysis

For testing RAP2.2 and RAP2.12 activities, promoters of *ADH1*, *PGB1* (see above) and *PDC1*, *SAD6* and *HRA1* (Bui et al., 2015) were placed in front of the firefly *LUC* gene in pGWL7,0 (Karimi et al., 2002). *35S:RAP2.2* or *35S:RAP2.12* in p2GW7,0 (Karimi et al., 2002) were co-transformed with reporter constructs and the normalisation vector containing *35S:RLUC* into protoplasts according to Wu et al. (2009) and Schmidt et al. (2013). For some assays, in addition to the mentioned plasmids, *35S:AtMED25* or *35S:OsMED25* or *35S:AtMED16* in p2GW7,0 was co-transformed. In case of SUB1A, *35S:SUB1A* in p35S-GAD-GW (Weltmeier et al., 2006) was co-transformed together with *pUAS:LUC* and either *35S:AtMED25* or *35S:OsMED25* (both in p2GW7,0). For each construct, 5 µg were used. Dual luciferase assays (Promega) with four independent replicates were employed as reported previously (Schmidt et al., 2013) and emitted luminescence signals were quantified using the SYNERGY MY (BioTek) plate-reader system.

To study interaction between AtMED25 and RAP2.2 or RAP2.12 in Arabidopsis mesophyll protoplasts, *35S:AtMED25* in pDH51-GW-YFPn was co-transformed with either *35S:RAP2.2* or *35S:RAP2.12* in pDH51-GW-YFPc following the tape-Arabidopsis-sandwich protocol (Wu et al., 2009). Complex formation of SUB1A with OsMED25 or AtMED25 was investigated by using the pE-SPYNE-GW and pE-SPYCE-GW vectors (Walter et al., 2004) in rice shoot protoplasts following the protocol of Schmidt et al. (2013). As negative controls, recombinant vectors were co-transformed with the corresponding counterpart containing GFP. For each construct, 5 µg were used.

### *In-vitro* pulldown assay

For the *in-vitro* pulldown assay, flag-tagged MED25 protein and CFP-tagged RAP2.2 and RAP2.12 proteins (Schmidt et al., 2018) were expressed using TNT SP6 High Yield Wheat Germ (Promega) following the manufacturer’s instructions. Subsequently, expression mixtures were filled up with 500 µl binding buffer (50 mM Tris-HCl, 150 mM NaCl) and 250 µl of MED25 samples was mixed with an equal volume of RAP2.2 or RAP2.12. The samples were placed on a rotator at 16 degrees for 2 hours to allow for complex formation. Subsequently, 25 µl calibrated magnetic anti-GFP beads were added (Chromotek), and the samples were incubated for an additional hour at 16 degrees. After this incubation, the beads were collected and washed 5 times with 500 µl washing buffer (50 mM Tris-HCl, 150 mM NaCl, 0.5% TritonX-100). Subsequently, beads were resuspended in laemmli buffer (with ß-ME) and heated to 95°C for 5 min to elute and denature the proteins. Eluted and input samples were analysed by 10% SDS-PAGE and immunoblotting with anti-FLAG antibody (Sigma) and anti-GFP antibody (Roche).

### Confocal microscopy

Imaging and analysis of GFP and YFP signals in order to detect protein localisation and interaction were performed using a LSM780 confocal microscope (Zeiss). Wavelengths for fluorophore excitation and emission were 488 nm and 500-550 nm for GFP and 514 nm and 520-550 nm for YFP respectively. Nuclei were stained using DAPI following the manufacturer’s advice and excitation and detection were obtained at 405 nm and 525-465 nm, respectively. Chlorophyll autofluorescence was detected at 655-685 nm. For AtMED25-GFP localisation in stable Arabidopsis plants, 2-week-old seedlings were analysed. The authors acknowledge the team of the Light Microscopy Technology Platform of the Faculty of Biology, Bielefeld University, Germany.

### RT-qPCR analysis

Extraction of RNA, DNA removal, cDNA synthesis and RT-qPCR analyses were performed as described previously (Schmidt et al., 2013). Four to five biological replicates were used for each experiment. UBIQUITIN10 served as reference gene according to Licausi et al. (2011). Calculation of relative transcript abundance was done using the comparative cycle threshold (CT) method (Livak and Schmittgen, 2001). Oligonucleotide sequences used for expression analysis of hypoxic genes can be found elsewhere (Schmidt et al., 2018). For ethylene, hypoxia treatments and subsequent RT-qPCR on *in vitro*-grown seedling root tips, protocols were used as described previously (Hartman et al., 2019), using *ADENINE PHOSPHORIBOSYL TRANSFERASE 1* (*APT1*) as a reference gene. Oligonucleotide sequences for RT-qPCR expression analysis related to hypoxia combined with ethylene can be found in **Supplemental Table S2.**

### RNA-SEQ analysis

Global transcript responses to hypoxia (3h, 1% oxygen) were analysed in 14-day-old *med25-1* and wildtype seedlings grown on horizontal half-strength MS plates containing 0.5% sucrose. For each genotype and treatment, 3 biological replicates were used. Extracted RNA was checked and processed as described before (Schmidt et al., 2018). RNA-seq data is deposited at NCBI’s Gene Expression Omnibus (GEO) (record number GSE130962).

### ChIP-qPCR analysis

ChIP experiments were essentially done as previously described (Kaufmann et al. 2010; Neuser et al., 2019). In brief, 0.7-0.8 g two-week-old *35S:AtMED25-GFP* seedlings were vacuum-infiltrated with MC buffer containing 1% formaldehyde. After nuclei isolation, the samples were sonicated to share the chromatin. Subsequently, immunopurification was performed using anti-GFP agarose beads (Chomotek) to enrich for MED25-GFP-bound DNA fragments. In addition, a negative control (binding control agarose beads, Chromotek) and input control were used. Immunoprecipitated chromatin was eluted three times with 100 μl of ice-cold glycine buffer [0.1 M glycine, 0.5 M NaCl, 0.05% Tween-20 (pH 2.8)]. Each eluate was buffered with 150 μl of Tris-HCl (pH 9.0) and subsequently pooled for each sample. After elution, samples were treated overnight with Proteinase K (Roche) at 37 °C. The next day, the proteinase K treatment was repeated for 6 hours at 65°C. The enriched DNA samples were precipitated overnight with 2.5 vol 100% ethanol, 1/10 of NaAc and 1 μl glycogen (Invitrogen) at −20 °C. Finally, samples were cleaned-up using the QIAquick PCR Purification Kit (Qiagen GmbH & Co. KG). Samples were eluted from the column in a 30 μl volume and analysed by RT-qPCR with specific primers (**Supplemental Table S3).**

RNAPII ChiP analysis was done similar as described above and previously (Hemsley et al., 2014), with a couple of modifications. A ChIP-grade antibody against Pol II C-terminal domain repeats (Ab5408; Abcam) was immobilised on MagnaChIP™ Protein A magnetic beads (Sigma-Aldrich). Both the wildtype as well as the *med25-1* mutant were used for the analysis under normoxic and hypoxic conditions. Primers used to amplify gDNA regions are provided in **Supplemental Table S3**).

### Gene ontology term enrichment analysis

Gene ontology (GO) term enrichment analysis was done using the agriGO v2.0 webtool (http://systemsbiology.cau.edu.cn/agriGOv2/) (Tian et al., 2017). GO terms with an FDR ≤ 0.05 were considered enriched. Heatmaps were generated using the Multiple Experiment Viewer (MeV) software (Saeed et al., 2017).

### Multiple sequence alignment

For the multiple sequence alignment of MED25 proteins from different species, sequences of homologous proteins were obtained from Phytozome v12.0 and aligned with Clustal X.

### Analysis of protein–protein interactions by IP-MS

GFP pull-down was performed as previously described by (Wendrich et al., 2017). Approximately 4 g of leaf material were harvested from each condition and genotype and flash frozen. Grinded material was homogenised in extraction buffer containing 50-mM Tris–HCl, pH 7.5, 0.15-M NaCl, 1 Protease inhibitor tablet/50 mL (Roche), and 1% (v/v) NP40). Protein extracts were sonicated, diluted (NP40 diluted to 0.2%) and then centrifuged twice for 15 min at 18000 rpm. After incubation for 2 h at 4°C, with anti-GFP-coated microbeads (μMACS, Miltenyi), protein extracts were applied on the μColumns (μMACS, Miltenyi), washed and then eluted. The elute was reduced with DTT, alkylated with iodocetamide and digested with sequencing grade Trypsin. The next day, the peptides were purified using the Bond Elut OMIX tips (Agilent). The sample concentration was determined, and an equal amount of peptides (600 ng) was injected and analysed by liquid chromatography–tandem mass spectrometry (LC–MS/MS) with a tandem UltiMate 3000 RSLCnano system in-line connected to an Orbitrap Fusion Lumos Tribrid mass spectrometer (Thermo Fisher Scientific). Raw files were processed with the MaxQuant software (1.6.9.0) as reported in He et al., 2021. ‘ProteinGroups’ file from MaxQuant was used as input in an R-script, data were filtered, Label Free Quantification (LFQ) intensities were LOG10 transformed and the lowest value is set as a detection threshold. Statistical T-TEST is performed and ratios and *p-*values are calculated (**Supplementaly Dataset S3**).

## Author contribution

J.S., J.vD. and R.S-S. designed the research. K.vB., L.L., S. Frohn, S. Frings, T.R., F.A., K.W.,

K.S., S.H., A.R., R.S., D.V., and A.M. carried out experiments. G.B., F.vB., S.H., J.vD., J.S., and R.S-S. analysed and visualised the data. JS, AM and RS-S wrote the manuscript, all authors edited the manuscript.

## Accession numbers

Sequence information from this article can be found in the Arabidopsis Genome Initiative database under the following accession numbers: *ADH1* (AT1G77120), *AEL1* (AT2G25760), *AEL3* (AT3G03940), *AtMED11* (AT3G01435), *AtMED12* (AT4G00450), *AtMED13* (AT1G55325), *AtMED16* (AT4G04920), *AtMED18* (AT2G22370), *AtMED19* (AT5G12230), *AtMED21* (AT4G04780), *AtMED22A* (AT1G16430), *AtMED25* (At1g25540), *AtMED28* (AT3G52860), *AtMED3* (AT3G09180), *AtMED30* (AT5G63480), *AtMED32* (AT1G11760), *AtMED34* (AT1G31360), *AtMED36* (AT4G25630), *AtMED8* (AT2G03070), *AtMED9* (AT1G55080), *APT1* (AT1G27450), *ATRBP45C* (AT4G27000), *C3H44* (AT3G51120), *ELKS1* (AT1G03290), *ELKS2* (AT4G02880), *ETR2* (AT3G23150), *HRA1* (AT3G10040), *HRE1* (AT1G72360), *HRE2* (AT2G47520), *HUP9* (AT5G10040), *J3* (AT3G44110), *LBD41* (AT3G02550), *MBR1* (AT2G15530), *MBR2* (AT4G34040), *OsERF66* (LOC_Os03g22170), *OsERF67* (LOC_Os07g47790), *OsMED25* (LOC_Os09g13610), *SUB1A* (DQ011598), *PDC1* (AT4G33070), *PCO1* (AT3G10040), *PCO2* (AT5G39890), *PGB1* (AT2G16060), *RAF20* (AT1G79570), *RAP2.12* (AT1G53910), *RAP2.2* (AT3G14230), *RAP2.3* (AT3G16770), *SAD6* (AT1G43800), *SUS1* (AT5G20830), *SUS4* (AT3G43190), *TOL6* (AT2G38410), *UBI10* (AT4G05320).

## Supplemental Data

**Supplemental Figure S1.** Yeast-two-hybrid assays for testing interaction between Mediator complex subunits with the ERFVII factors RAP2.2 and RAP2.12.

**Supplementaly Figure S2.** RNA-Seq data for hypoxia core genes in hypoxia-treated *med25-1* and wildtype.

**Supplemental Figure S3.** Position of primer pairs used in RNAPII-specific ChIP assay.

**Supplemental Figure S4.** Yeast-two-hybrid assays for testing interaction between Mediator complex *At*MED25 with other Mediator subunits from Arabidopsis.

**Supplemental Figure S5.** Multiple sequence alignment of MED25 proteins from different plant species.

**Supplemental Table S1.** Oligonucleotide sequences used for cloning constructs in pENTR-D/TOPO and other vectors.

**Supplemental Table S2.** Sequences of oligonucleotides used for RT-qPCR expression analysis of ethylene-induced genes.

**Supplemental Table S3.** Sequences and positions of motifs and oligonucleotides used for ChIP-qPCR analysis.

**Supplemental Dataset S1:** Transcriptome and GO term analysis of wild-type and *med25-1* plants under control and hypoxia.

**Supplemental Dataset S2.** GO term analysis of specific and overlapping DEGs in wildtype and *med25-1* upon hypoxia.

**Supplemental Dataset S3.** List of interactors of *At*MED25 identified by GFP pull-down experiments in Arabidopsis rosettes.

## Acknowledgement

We thank Alina Beling, Brigitta Ehrt, Patricia Dalcin Martins and Christian Hammerschmid for technical support. We furthermore acknowledge Rens Voesenek for his valuable intellectual input. Seeds of *med25-2* were kindly provided by Yunhai Li (Institute of Genetics and Developmental Biology, Chines Academy of Sciences, Beijing, China) while *erfvii* seeds were a gift from Michael Holdsworth (School of Biosciences, University of Nottingham, UK). We thank Li-Jia Qu (School of Life Sciences, Peking University, Beijing, Peking) for providing *med16* seeds and *AtMED8*, *AtMED16*, *AtMED19*, *AtMED21*, *AtMED28*, *AtMED32*, *AtMED34* and *AtMED36* coding sequences in pENTR. We further thank Francesco Licausi (University of Oxford, UK) for giving *PDC1*, *SAD6* and *HRA1* promoters in pGWL7,0 and Beatrice Giuntoli (Dipartimento di Biologia, Università di Pisa, Italy) for providing split-YFP empty vectors. We acknowledge the kind providing of *HRE1*, *HRE2* and *RAP2.3* coding sequences in pENTR by Margret Sauter (Christian-Albrechts University, Kiel, Germany). We are further grateful to Pamela C. Ronald (Department Plant Pathology and the Genome Center, University of California, Davis, USA) for providing *SUB1A* coding sequence in pENTR.

