## Supplementary material for "MEDIATOR SUBUNIT 25 modulates ERFVII-controlled hypoxia responses in Arabidopsis": Supplemenal Figures and Tables

**Supplementary Figures and Tables – Schippers et al. Plant Cell**

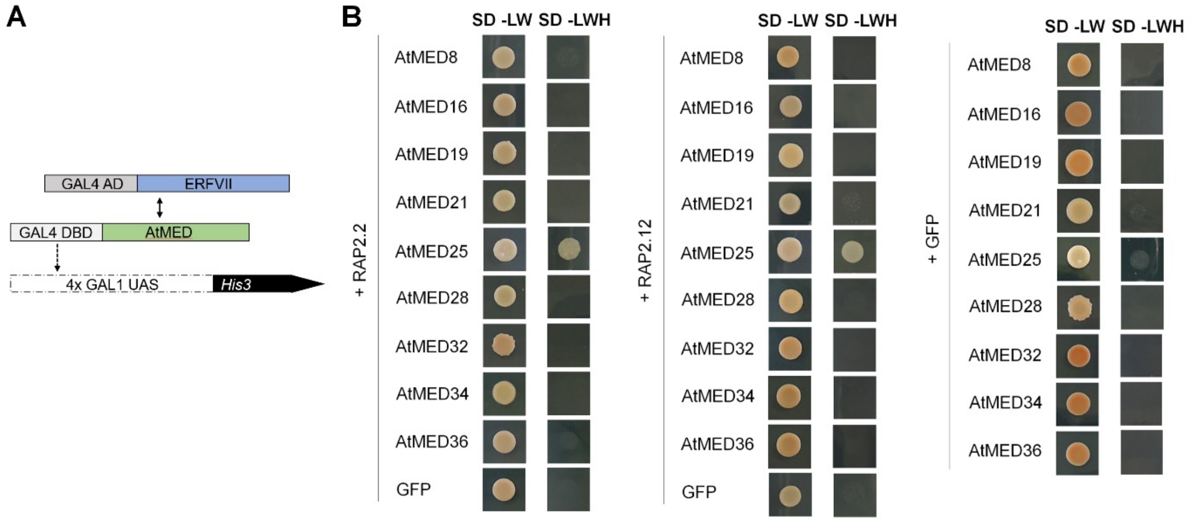


**Supplementary Figure S1.** Yeast-two-hybrid assays for testing interaction between Mediator complex subunits with the ERFVII factors RAP2.2 and RAP2.12. (Supports Figure 1)

(**A**) Schematic representation of yeast-two-hybrid assay performed with ERFVIIs and AtMED subunits. (**B**) Growth of yeast colonies on selection medium (-LWH) lacking leucine (L), tryptophan (W), and histidine (H) with 65 mM 3-aminotriazole (3-AT) indicates interaction.

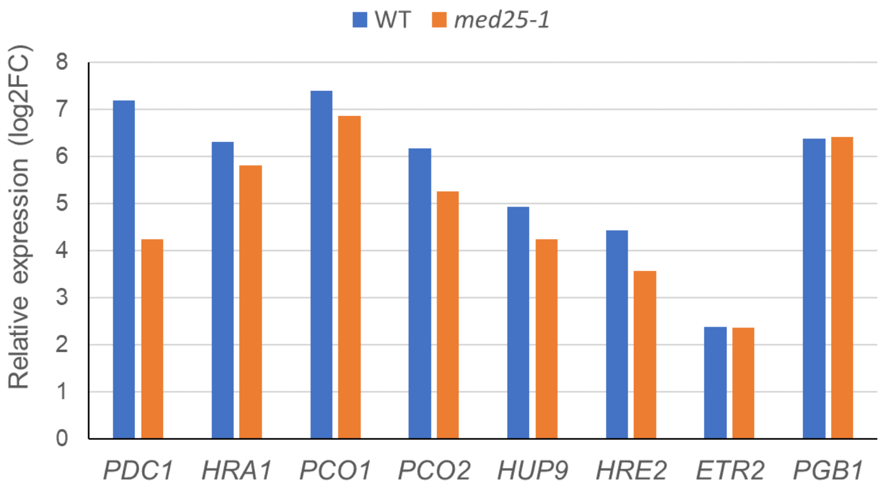


**Supplementary Figure S2.** Plot of RNA-Seq data for selected hypoxia core genes in hypoxia-treated *med25-1* and wildtype. (Supports Figure 2).

Plotted are the $log_2FC$ (FC, fold change) values for selected hypoxia core genes as reported in Supplemental Dataset 1.

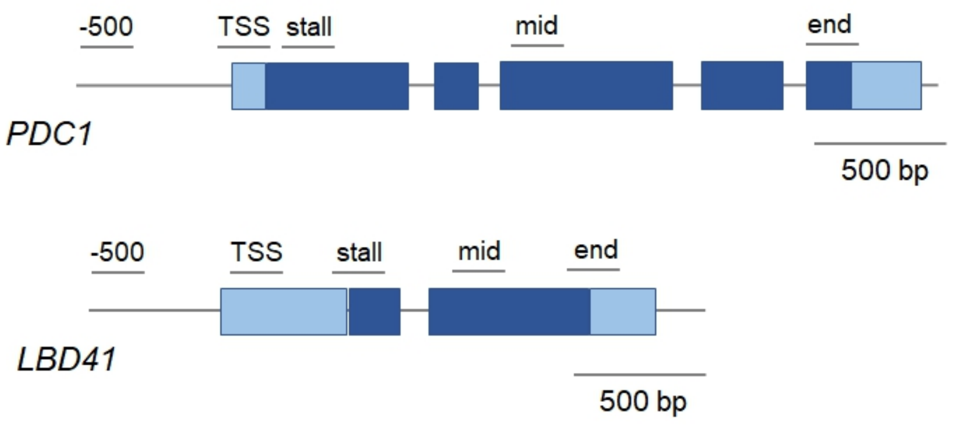


**Supplementary Figure S3.** Position of primer pairs used in RNA Pol II ChIP assay. (Supports Figure 3).

Overview of primer pair locations along two hypoxia-responsive genes. Dark blue boxes indicate the exons of the genes, while light blue boxes represent the UTRs. Thin black lines above show the regions amplified by the primers in each ChIP reaction.

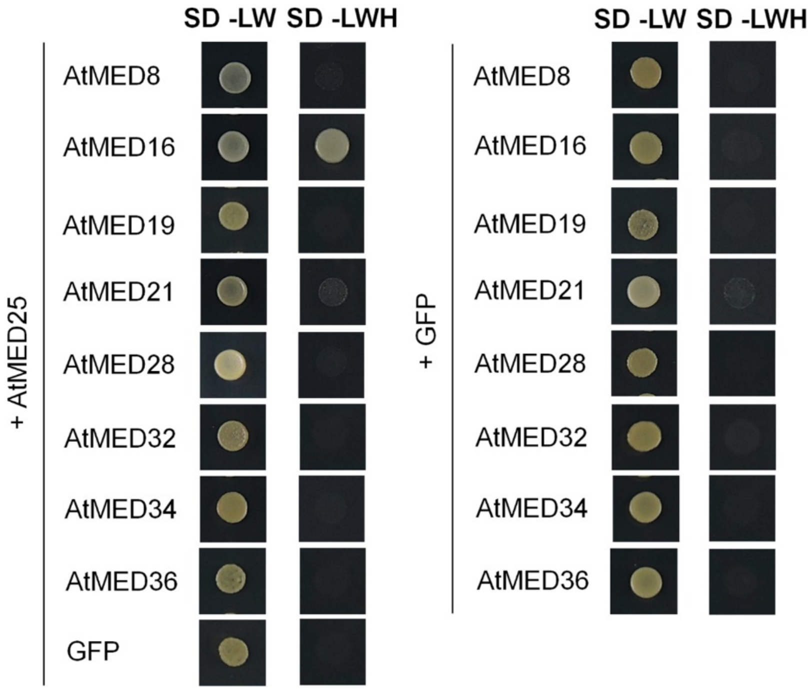


**Supplementary Figure S4.** Yeast-two-hybrid assays for testing interaction between Mediator complex *At*MED25 with other Mediator subunits from Arabidopsis. (Supports Figure 5)

Growth of yeast colonies on selection medium (-LWH) lacking leucine (L), tryptophan (W), and histidine (H) with 65 mM 3-aminotriazole (3-AT) indicates interaction.

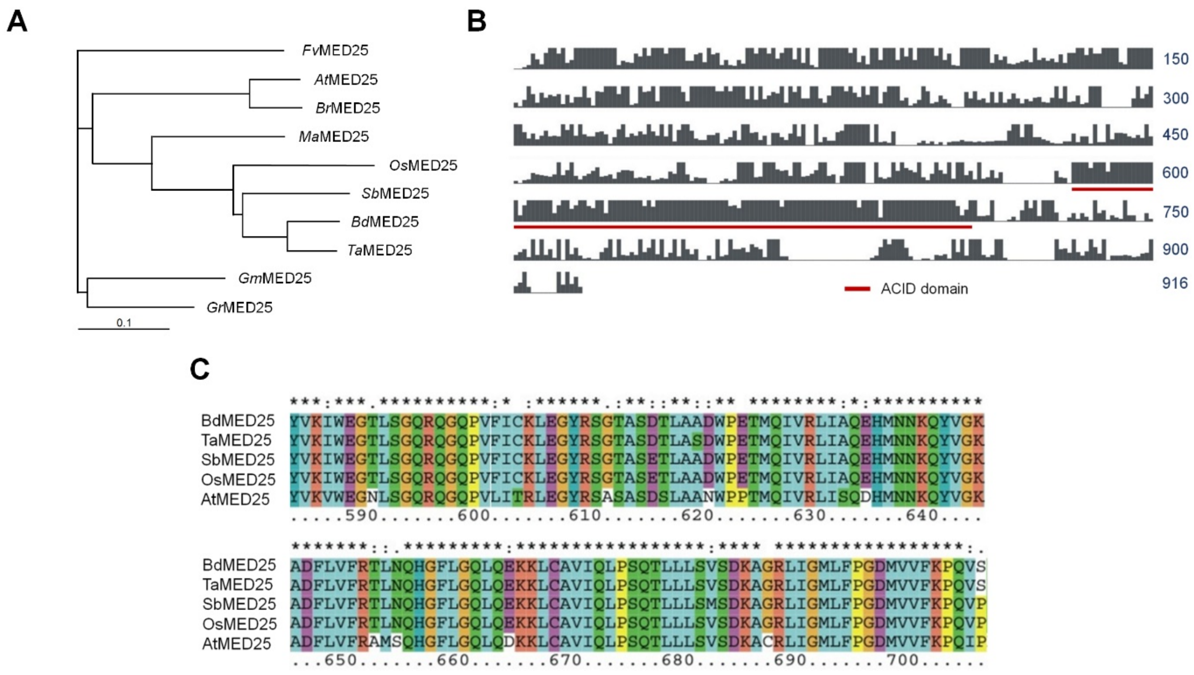


**Supplementary Figure S5.** Multiple sequence alignment of MED25 proteins from different plant species. (Supports Figure 6)

(**A**) Phylogenetic tree of *At*MED25 and its homologous proteins from selected species. Dendrogram obtained using neighbor-joining analysis. (**B**) Schematic representation of the sequence alignment of *At*MED25 and homologous proteins. Black boxes indicate conserved residues and the red underlined domain represents the activator-interacting domain (ACID). (**C**) Multiple sequence alignment indicates a strong conservation of the ACID of MED25 from *Brachypodium distachyon* (*Bd*), *Triticum aestivum* (*Ta*), *Sorghum bicolor* (*Sb*), *Oryza sativa* (*Os*) and *Arabidopsis thaliana* (*At*).

**Supplementary Table 1. Oligonucleotide sequences used for cloning constructs in pENTR-D/TOPO and other vectors.**

| Constructs | Length (aa) | Primer name | 5'→3' sequence | Usage |
|---|---|---|---|---|
| RAP2.2 1-379 (stop codon) | 1-379 | RAP2.2_FL_F | caccATGTGTGGAGGAGCTATAATCT | Y2H, transactivation assay |
| | | RAP2.2_FL_stop_R | TCAAAAGTCTCCTTCCAGCATGA | |
| RAP2.2 1-379 (no stop codon) | 1-379 | RAP2.2_Fl_F | caccATGTGTGGAGGAGCTATAATCT | BiFC |
| | | RAP2.2_FL_nostop_R | AAAGTCTCCTTCCAGCATGAAA | |
| RAP2.2_1-356 (stop codon) | 1-356 | RAP2.2_FL_F | caccATGTGTGGAGGAGCTATAATCT | Y2H, transactivation assay |
| | | RAP2.2_1-356_stop_R | TTATTCCTCTTCCTGAGTCACAGC | |
| RAP2.2_1-331 (stop codon) | 1-331 | RAP2.2_FL_F | caccATGTGTGGAGGAGCTATAATCT | Y2H, transactivation assay |
| | | RAP2.2_1-331_stop_R | TTAGTCAAGGTATGCCATCAGATCGTC | |
| RAP2.2_267-379 (stop codon) | 267-379 | RAP2.2_267-379_F | caccATGGGATACCAGTATTTCAGTTCCGA | Y2H, transactivation assay |
| | | RAP2.2_267-379_stop_R | TCAAAAGTCTCCTTCCAGCATGAAA | |
| RAP2.2_44-303 (stop codon) | 44-303 | RAP2.2_44-303_F | caccATGGATTTCTTCGATCTTGACGATGATT | Y2H |
| | | RAP2.2_44-303_stop_R | TCAATTGACAAGCATTGAAGAGATCTC | |
| RAP2.2_44-267 (stop codon) | 44-267 | RAP2.2_44-303_F | caccATGGATTTCTTCGATCTTGACGATGATT | Y2H |
| | | RAP2.2_44-267_stop_R | TCATCCATTGTTACCTCCAGCATCGAA | |
| RAP2.2_90-303 (stop codon) | 90-303 | RAP2.2_90-303_F | caccATGGCTTCCGCTTTCGTCTCCACT | Y2H |
| | | RAP2.2_44-303_stop_R | TCAATTGACAAGCATTGAAGAGATCTC | |
| RAP2.2_90-267 (stop codon) | 90-267 | RAP2.2_90-267_F | caccATGGCTTCCGCTTTCGTCTCCACT | Y2H |
| | | RAP2.2_44-267_stop_R | TCATCCATTGTTACCTCCAGCATCGAA | |
| RAP2.2_336-379 (stop codon) | 336-379 | RAP2.2_336-379_F | caccATGGCCTTGTGGGACACCCCACT | Y2H, transactivation assay |
| | | RAP2.2_FL_stop_R | TCAAAAGTCTCCTTCCAGCATGA | |
| RAP2.2_355-379 (stop codon) | 355-379 | RAP2.2_355-379_F | caccATGGTGACTCAGGAAGAGGAAAAC | Y2H, transactivation assay |
| | | RAP2.2_FL_stop_R | TCAAAAGTCTCCTTCCAGCATGA | |
| RAP2.12 1-358 (stop codon) | 1-358 | RAP2.12_FL_F | caccATGTGTGGAGGAGCTATAATATC | Y2H, transactivation assay |
| | | RAP2.12_FL_stop_R | TCAGAAGACTCCTCCAATCATG | |
| RAP2.12 1-358 (no stop codon) | 1-358 | RAP2.12_FL_F | caccATGTGTGGAGGAGCTATAATATC | BiFC |
| | | RAP2.12_FL_nostop_R | GAAGACTCCTCCAATCATG | |
| RAP2.12_1-339 (stop codon) | 1-339 | RAP2.12_FL_F | caccATGTGTGGAGGAGCTATAATATC | Y2H, transactivation assay |
| | | RAP2.12_1-339_stop_R | TTAGTTTGCACCATTGTCCTGAGTC | |
| RAP2.12_1-307 (stop codon) | 1-307 | RAP2.12_FL_F | caccATGTGTGGAGGAGCTATAATATC | Y2H, transactivation assay |
| | | RAP2.12_1-307_stop_R | TTACTTGAGCTTCTTAGCTGGATTGGC | |
| | 44-289 | RAP2.12_44-289_F | caccATGAATTTCTTCGATTTTGACGCTGAG | Y2H |

| | | | | |
|---|---|---|---|---|
| RAP2.12_44-289 (stop codon) | | RAP2.12_44-289_stop_R | TCAGTTGATAACCGCAGAAGAGATGT | |
| RAP2.12_44-253 (stop codon) | 44-253 | RAP2.12_44-253_F | caccATGAATTTCTTCGATTTTGACGCTGAG | Y2H |
| | | RAP2.12_44-253_stop_R | TCACCCATTACATCCAGCATCAACGGA | |
| RAP2.12_93-289 (stop codon) | 93-289 | RAP2.12_93-289_F | caccATGGTCTCCGCCGCTGCGGAAGGTT | Y2H |
| | | RAP2.12_44-289_stop_R | TCAGTTGATAACCGCAGAAGAGATGT | |
| RAP2.12_93-253 (stop codon) | 93-253 | RAP2.12_93-253_F | caccATGGTCTCCGCCGCTGCGGAAGGTT | Y2H |
| | | RAP2.12_44-253_stop_R | TCACCCATTACATCCAGCATCAACGGA | |
| RAP2.12_1-179 (stop codon) | 1-179 | RAP2.12_1-179_F | caccTGTGTGGAGGAGCTATAATATC | Y2H, transactivation assay |
| | | RAP2.12_1-179_stop_R | TTAATTCACCTTAGCTTTAGATCCA | |
| RAP2.12_180-358 (stop codon) | 180-358 | RAP2.12_180-358_F | caccATGTTCCCTGAAGAAAACATGAAGGCT | Y2H, transactivation assay |
| | | RAP2.12_FL_stop_R | TCAGAAGACTCCTCCAATCATG | |
| RAP2.12_225-358 (stop codon) | 225-358 | RAP2.12_225-358_F | caccATGTGTTTCATGGAGGAGAAACACCA | Y2H, transactivation assay |
| | | RAP2.12_FL_stop_R | TCAGAAGACTCCTCCAATCATG | |
| RAP2.12_255-358 (stop codon) | 255-358 | RAP2.12_255-358_F | caccATGCAGTATTTCAGCTCTGACCAGG | Y2H, transactivation assay |
| | | RAP2.12_FL_stop_R | TCAGAAGACTCCTCCAATCATG | |
| RAP2.12_309-358 (stop codon) | 309-358 | RAP2.12_309-358_F | caccATGATGGATTTCGAGACACCTTACAA | Y2H, transactivation assay |
| | | RAP2.12_FL_stop_R | TCAGAAGACTCCTCCAATCATG | |
| RAP2.12_335-358 (stop codon) | 335-358 | RAP2.12_335-358_F | caccATGCAGGACAATGGTGCAAACCCT | Y2H, transactivation assay |
| | | RAP2.12_FL_stop_R | TCAGAAGACTCCTCCAATCATG | |
| RAP2.12_1-23 (stop codon) | 1-23 | RAP2.12_FL_F | caccATGTGTGGAGGAGCTATAATATC | Y2H |
| | | RAP2.12_1-23_stop_R | CTAAAACTCGCTAGTAAC | |
| RAP2.12_1-79 (stop codon) | 1-79 | RAP2.12_FL_F | caccATGTGTGGAGGAGCTATAATATC | Y2H |
| | | RAP2.12_1-79_stop_R | CTAATCGGCGAAAACATC | |
| RAP2.12_1-103 (stop codon) | 1-103 | RAP2.12_FL_F | caccATGTGTGGAGGAGCTATAATATC | Y2H |
| | | RAP2.12_1-103_stop_R | CTAACCAAAAACTGAACC | |
| RAP2.12_1-123 (stop codon) | 1-123 | RAP2.12_FL_F | caccATGTGTGGAGGAGCTATAATATC | Y2H |
| | | RAP2.12_1-123_stop_R | CTAATTCTTCCTCTTCCTATT | |
| AtMED25 full length (stop codon) | | AtMED25_FL_F | caccATGTCGTCGGAGGTGAAACAG | Y2H, transactivation assay |
| | | AtMED25_FL_stop_R | TTATCCCATGAAGCCAGCTCC | |
| AtMED25 full length (no stop codon) | | AtMED25_FL_F | caccATGTCGTCGGAGGTGAAACAG | GFP fusion, in planta localization, BiFC, ChIP |
| | | AtMED25_FL_nostop_R | TCCCATGAAGCCAGCTCCAG | |
| AtMED25 promoter | | pAtMED25_F | caccTTGTTTAGATTTATTCGGATTTTA | GUS analysis |
| | | pAtMED25_R | AGAAAATGGGATATACAAAGGA | |

| | | | | |
|---|---|---|---|---|
| ADH1 promoter | | pADH1_F | caccTGGGCCTATGATTCAACACAACA | Transactivation assay |
| | | pADH1_R | TATCAACAGTGAAGAACTTGCTT | |
| HB1 promoter | | pHB1_F | caccGAAATTAAACACCTTCTTCTGTAA | Transactivation assay |
| | | pHB1_R | AATATTTCACAACCTCTAAATGAT | |
| OsMED25 full length (stop codon) | | OsMED25_FL_F | caccATGGCGGCGGCGGCGGCCGAGA | Y2H, transactivation assay, BiFC |
| | | OsMED25_FL_stop_R | TCAAGATAGGTAGCCACCCCCA | |
| RAP2.2-GFP | | RAP2.2-CFP_F | TTTGCGATCGCATGTGTGGAGGAGCTATAATCT | Co-IP |
| | | RAP2.2-CFP_R | AAAGTTTAAACTTACTTGTACAGCTCGTCCATGC | |
| flag-AtMED25Δq | | flag-AtMED25Δq_F | AAAGCGATCGCATGGATTACAAGGATGACGATGACAAGGCAGCCGGTTCGTCGGAGGTGAAACAGCTGA | Co-IP |
| | | flag-AtMED25Δq_F | TGTGGTTTAAATTACACAACCATATCCCCTGGGAAAAGCATTCCAA | |

**Supplementary Table 2. Sequences of oligonucleotides used for RT-qPCR expression analysis of ethylene-induced genes.**

| **Gene** | **Forward primer 5'→3'** | **Reverse primer 5'→3'** |
|---|---|---|
| *ETR2* | tgttagattctccggcggctatg | ttcccatgaatcaactgcaccac |
| *PGB1* | ggctcttgtagtgaagtcttgga | cttcgttgttggtgcaatctca |
| *HRE1* | tccgatgagccatttgtcttctcc | ccatcttccccaaggccttc |
| *RAP2.2* | cctagcgtcgtatcccagaa | agggtttgcaccattgtcctgag |
| *RAP2.3* | aactcacggctgaggaactctg | acgttaacttggttggtgggatgg |

**Supplementary Table S3. Sequences and positions of motifs and oligonucleotides used for ChIP-qPCR analysis.**

| Promoter | Protein | Motif | Motif sequence 5'→3' | Motif position 5'→3' | Forward primer 5'→3' | Reverse primer 5'→3' |
|---|---|---|---|---|---|---|
| *PDC1* | *At*MED25-GFP | R1 | GCAGATGGTTTT | -685 till - 674 | aagaagcaaaccccaaaacc | gccatttgcatttggtcag |
| *PDC1* | *At*MED25-GFP | R2 | GGGAGAGGTTTT | -125 till - 114 | gaatgtttgggtcaaatctcaa | aacaaaatgagaatgggagagg |
| *LBD41* | *At*MED25-GFP | R1 | GCCGACGCTTTC | -898 till - 887 | aaagcgtcggctaacagaga | tccggtttttgatcttttcttc |
| *LBD41* | *At*MED25-GFP | R2 | GCTCCTGGTTTT | -748 till - 736 | tttccagaaaaccaggagcta | tgggatttgagtcatttgagg |
| *LBD41* | *At*MED25-GFP | R3 | GCCGCTGTTTTT | -349 till - 338 | cattggatagagtggggacaa | ccctgattcaaggttttgct |
| *SAD6* | *At*MED25-GFP | R1 | none | none | tggcattctcgtgtacaaagtc | tgtgatcttgctgttcaattgtt |
| *SAD6* | *At*MED25-GFP | R2 | CCCCACGGTTTG | -153 till - 142 | cccacagaatctcaaaccag | tggttggctgttgttcgtta |
| *UBI10* | *At*MED25-GFP | R1 | none | none | aaccactttgacgccgttta | acggctggatcttatgacga |
| *LBD16** | *At*MED25-GFP | R1 | TGTCTC | -989 till - 984 | cccaataaattagaagtctcatgttgc | caaatttatcgagtgagccaaag g |
| *PDC1* | RNAPII | up500 | none | none | accccaaaaccatctgcatct | caagagacgtgatgccatttg |
| *PDC1* | RNAPII | TSS | none | none | tcacacacatacacaaacttgaca | gtcggcttgcaatcatcgatc |
| *PDC1* | RNAPII | stall | none | none | gatgattgcaagccgacgaac | gatggtgatagcggaggaagg |
| *PDC1* | RNAPII | mid | none | none | tcaatactctgtttctgcgaga | ccgcagcttctaaacccatct |
| *PDC1* | RNAPII | end | none | none | agaggagttagtggaggcgat | gccccactcaagcaactct |
| *LBD41* | RNAPII | up500 | none | none | gtgctacaaaatgggtctcaca | tccagacaaccaaaactcagct |
| *LBD41* | RNAPII | TSS | none | none | atgacacgcgcattggataga | gcttcttgtgtttcttcccca |
| *LBD41* | RNAPII | stall | none | none | tcggaaagggtgtagtgagga | ttcaggcgatttgatccaagc |
| *LBD41* | RNAPII | mid | none | none | gtggaggctgtgatgaaagga | cgtagatcttaagaggcggacc |

| | | | | | | |
|---|---|---|---|---|---|---|
| *LBD41* | RNAPII | end | none | none | agagcacgtgtaagactgagc | agttaccggctcgacttgaag |

* Ito et al., 2016
